# Energetic and structural control of polyspecificity in a multidrug transporter

**DOI:** 10.1101/2025.04.09.647630

**Authors:** Silas T. Miller, Katherine A. Henzler-Wildman, Srivatsan Raman

## Abstract

Multidrug efflux pumps are dynamic molecular machines that drive antibiotic resistance by harnessing ion gradients to export chemically diverse substrates. Despite their clinical importance, the molecular principles underlying multidrug promiscuity and energy efficiency remain poorly understood. Using multiparametric deep mutational scanning across eight substrates and two energy conditions, we deconvolute the contributions of substrate recognition, energetic coupling, and protein stability, providing an integrated, high-resolution view of multidrug transport. We find that substrate specificity arises from a distributed network of residues extending beyond the binding site, with mutations that reshape binding, coupling, conformational flexibility, and membrane interactions. Further, we apply a pH-based selection scheme to measure the effect of mutation on pH-dependent transport efficiency. By integrating these data, we reveal a fundamental relationship between efficiency and promiscuity: highly efficient variants exhibit broad substrate profiles, while inefficient variants are narrower. These findings establish a direct link between energy coupling and polyspecificity, uncovering the biochemical logic underlying multidrug transport.

## Introduction

Promiscuous membrane transporters, termed multidrug efflux pumps, contribute to multidrug resistance (MDR) in cancers and bacterial infections by exporting chemically diverse substrates^1,2^. The ability to efflux dissimilar substrates while remaining selective for toxins presents a fundamental biochemical paradox. In traditional models of ligand recognition, precisely arranged functional groups accommodate a single substrate or small set of structurally related molecules. In contrast, multidrug efflux pumps dynamically recognize and expel many structurally dissimilar molecules, challenging core principles of protein-ligand interactions and raising questions about the molecular basis of substrate recognition in transporters^3^.

Three basic challenges have hindered progress in understanding multidrug transport. First, membrane protein structures remain difficult to solve, limiting structural descriptions of polyspecificity. While cryo-electron microscopy (cryo-EM) has yielded structural insights into some multidrug transporters^4–7^, deriving functional detail from static structures is not straightforward, especially for transporters which undergo large (>10Å) conformational changes. Second, transport is mechanistically complex, involving energy coupling, conformational change, and dynamic substrate binding and release. Traditional mutational studies of polyspecificity primarily examine the substrate binding site^8–11^, but this focus overlooks the multidomain interactions that govern efflux, including the well-documented impact of distal mutations on gating and regulatory mechanisms^12–17^. Third, uncovering the basis of promiscuity requires profiling of many substrates, but conventional biochemical assays are too laborious to test both mutants and substrates at scale. Most studies characterize transporter mutants with only one or a few substrates, failing to capture the multidimensional nature of substrate recognition—the very basis of their clinical and biochemical significance.

Multidrug efflux pumps move substrates against a concentration gradient and therefore must consume energy to function. Proton-coupled efflux pumps harness energy from the proton motive force (PMF). Like specificity, the mechanistic details of energy coupling are flexible and not well understood^18–25^. A key question is whether broad specificity imposes an energetic cost. One hypothesis suggests that the structural flexibility required for polyspecificity compromises overall transport efficiency^26,27^. Another perspective argues that efficiency may actually enable promiscuity: since substrates with varying enthalpies of binding impose different energetic demands, an optimally efficient transporter may be better equipped to accommodate a diverse range of molecules. While this interplay between energy efficiency and polyspecificity has been explored in ATP-driven transporters on a limited scale^28^, a comprehensive understanding remains elusive.

Together, these challenges highlight a fundamental gap in our understanding of multidrug efflux pumps: the molecular basis of polyspecificity and the tradeoffs between promiscuity and efficiency remain unsolved. Resolving these questions is essential for understanding efflux-mediated resistance and could inform new strategies to combat MDR in the clinic.

To dissect the molecular basis of polyspecificity and energy efficiency in multidrug transporters, we systematically analyzed the NorA efflux pump using deep mutational scanning (DMS) across eight diverse substrates and two proton gradient conditions. NorA, a proton-coupled efflux pump of *Staphylococcus aureus*, confers resistance to fluoroquinolones, antiseptics, disinfectants, and other antibiotics^29–31^, and is frequently overexpressed in MDR infections, including methicillin-resistant *S. aureus* (MRSA).^32^ NorA’s broad substrate profile, clinical relevance, and role in human disease—contributing to nearly 100,000 deaths in 2019^33^—make it an ideal model to uncover the principles of multidrug transport. Because DMS requires that function be linked to a selectable phenotype like viability—collapsing mechanistic nuance into a single measurement— we developed multiparametric screens to deconvolute functional effects into distinct elements of fitness: transport (drug resistance), energy efficiency (pH sensitivity), and stability (Rosetta ΔΔG).

Our data reveal that substrate specificity is not confined to the binding site but arises from a distributed network of residues throughout the protein. Mutations at these sites reshape energy coupling, conformational dynamics, membrane interactions, and substrate binding, collectively tuning transport activity. Detailed analysis of substrate-specific variants shows that subtle sequence changes can predictably reprogram substrate preference, highlighting remarkable adaptability encoded within the transporter architecture. By measuring pH-dependent transport efficiency as a proxy for energy coupling, we reveal a fundamental coupling between energy utilization and substrate breadth: efficient variants maintain broad specificity, while inefficient variants have restricted substrate profiles. Finally, we develop a thermodynamic framework that links energy efficiency to substrate promiscuity, uncovering the biochemical logic by which multidrug transporters achieve broad substrate recognition while maintaining energy efficiency. Together, these findings establish fundamental principles of transporter polyspecificity and provide a blueprint for understanding, predicting, and engineering substrate selectivity in these dynamic molecular machines.

## Results

### Deep mutational scanning of NorA maps sequence to function for eight substrates

NorA is a proton-coupled multidrug transporter of *S. aureus* that plays a key role in drug resistance by exporting a variety of clinically relevant antimicrobials. Recent cryo-EM structures of NorA have described its proton-coupled transport mechanism^4,34,35^. As a member of the Major Facilitator Superfamily (MFS), NorA adopts the characteristic 12-transmembrane (TM) helix fold, organized into two 6-TM bundles^36^. Within the binding site, four charged residues—Glu222, Asp307, Arg98, and Arg310—participate in proton translocation^4,34,37^ (Fig. 1A). In the outward-open conformation, Glu222 and Asp307 bind protons, triggering conformational change where the N– and C-terminal bundles rotate to the inward-open conformation, exposing the binding site to the cytosol. Protons are then released, allowing the antimicrobial substrate to bind. This cycle enables NorA to harness the inward-directed PMF to drive the efflux of toxic compounds.

**Figure 1.**
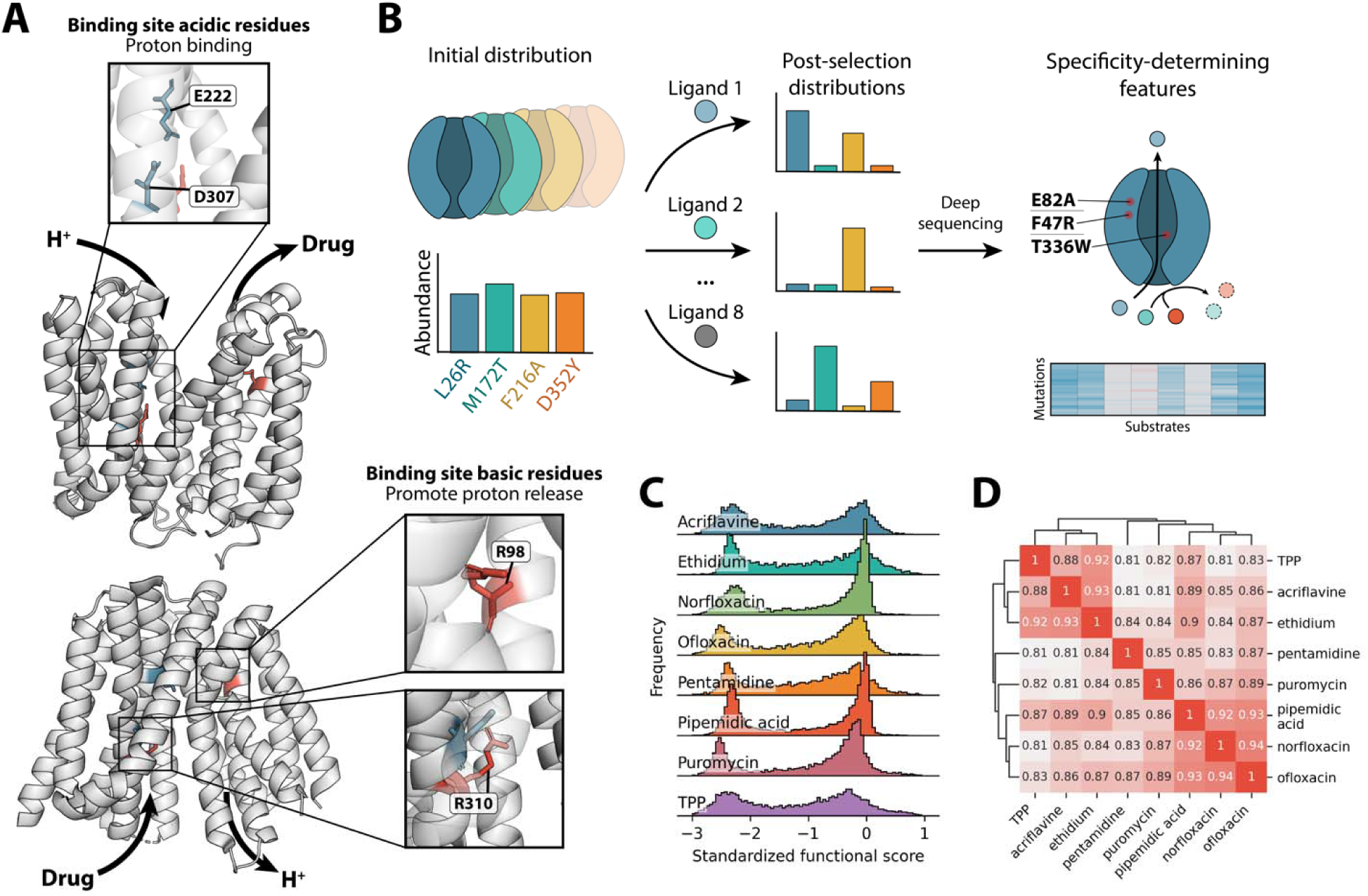
Deep mutational scanning identifies substrate-specific functional landscapes of the multidrug efflux pump NorA. [A] Structures of NorA in the outward-open (7LO8, top) and inward-open (9B3M, bottom) conformations. Key residues involved in proton-coupling are highlighted. [B] Deep mutational scanning using multiple substrates for selective pressure will reveal features governing specificity. [C] Distributions of variant functional scores across the eight substrates tested. [D] Clustered pairwise Spearman correlations between functional score profiles obtained with each substrate.

To systematically dissect polyspecificity in NorA, we applied deep mutational scanning (DMS)— a high-throughput approach that maps the functional impact of mutations across the entire protein sequence^38^. We introduced a pooled single-amino acid substitution library of NorA variants into bacterial populations and subjected them to selection with a toxic NorA substrate. We used deep sequencing to measure changes in variant abundance before and after selection. Variants that export the substrate and confer resistance allow cells to survive and proliferate, while loss-of-function variants are depleted during selection.

To investigate the basis of polyspecificity, we performed parallel DMS screens with eight substrates, generating distinct sequence-function landscapes for each (Fig. 1B). This approach identifies mutations with substrate-dependent effects on function, as well as residues essential for transport of all substrates. By integrating these data, we aim to uncover the molecular rules of substrate recognition, specificity, and the energetic constraints of multidrug efflux.

DMS typically focuses on small proteins or domains that can be directly sequenced with short-read Illumina sequencing (500–600bp). Because NorA exceeds this range at ∼1.2kb, we incorporated 20-nucleotide random DNA barcodes to uniquely identify each variant. Barcode-variant relationships were determined with short-read sequencing via separate “mapping constructs” which position variants adjacent to their barcode (Supplementary Fig. 1A). Barcodes were filtered for quality and empirically confirmed using a sub-library spanning 246bp (Supplementary Fig. 1B-E; see Methods). The pre-selection library contained 99.7% of possible missense and nonsense mutations (388 aa x 20 mut/aa = 7,760 possible variants), distributed relatively evenly across all positions with 31 barcodes per variant on average (Supplementary Fig. 2A-C). The library was expressed in *E. coli* deleted of its primary efflux machinery (MG1655-acrAB) to limit confounding effects of the host’s native efflux pumps. This genotype is a well-established background for studying multidrug efflux in *E. coli* and is hypersensitive to most antimicrobials without growth deficiency absent selection^39–42^.

We obtained functional scores for each mutant in the library (N = 7,730) in the context of eight antimicrobial substrates: acriflavine, ethidium, norfloxacin, ofloxacin, pentamidine, pipemidic acid, puromycin, and tetraphenylphosphonium (TPP) (Supplementary Fig. 3-10) (Total 7,730 variants/substrate x 8 substrates = 61,840 variant-function datapoints). These substrates were chosen to maximize clinical relevance and structural diversity, representing a range of drug classes, hydrophobicities, sizes, polar surface areas, and charges (Supplementary Table 1).

The distribution of functional scores is bimodal (Fig. 1C), with a peak around zero (wild type-like activity) and –2.5 (complete loss of activity). This is consistent with the expectation that most mutations either have no effect or abolish activity altogether. Each distribution also has a small positive tail, representing variants that outperform wild type. Many gain-of-function variants recur across all eight selections, suggesting a substrate-independent enhancement such as improved energy efficiency. While the distributions have high-level similarities across substrates, they differ in active/inactive peak sizes and prevalence of variants with intermediate activity, indicating substrate-specific effects.

To evaluate whether our dataset captured meaningful biochemical relationships between substrates, we computed pairwise Spearman correlations of functional profiles for all tested compounds. Correlation coefficients ranged from 0.81 to 0.94 (Fig. 1D), with the highest values observed among chemically similar substrates. For example, the polyaromatic cations acriflavine, ethidium, and TPP showed strong mutual correlations (0.88–0.93), as did the (fluoro)quinolone antibiotics norfloxacin, ofloxacin, and pipemidic acid (0.92–0.94). In contrast, puromycin and pentamidine—which share few similarities with the other substrates—exhibited lower correlations.

The pattern of correlations, which aligns with substrate chemical similarity, suggests that distinct sequence features mediate transport of different substrate classes, and gives confidence that our DMS screen accurately reflects the underlying biochemistry of NorA-mediated efflux.

### Unsupervised hierarchical clustering reveals sequence determinants of specificity

NorA’s overall specificity code is embedded in the unique substrate profile of each variant. To uncover this code without bias toward specific regions or mutations, we used unsupervised hierarchical clustering to group variants by their activity profiles (Supplementary Fig. 11). This revealed three major groups: universally permitted mutations (cluster of N = 3,392 variants), which have wild type–like activity across all substrates; universally disabling mutations (cluster of N = 1,912 variants), inactive on all substrates; and mutations with substrate-dependent effects (28 clusters ranging from N = 3 to N = 58 variants) (Fig. 2A). We expect that this third group encodes the molecular basis of polyspecificity.

**Figure 2.**
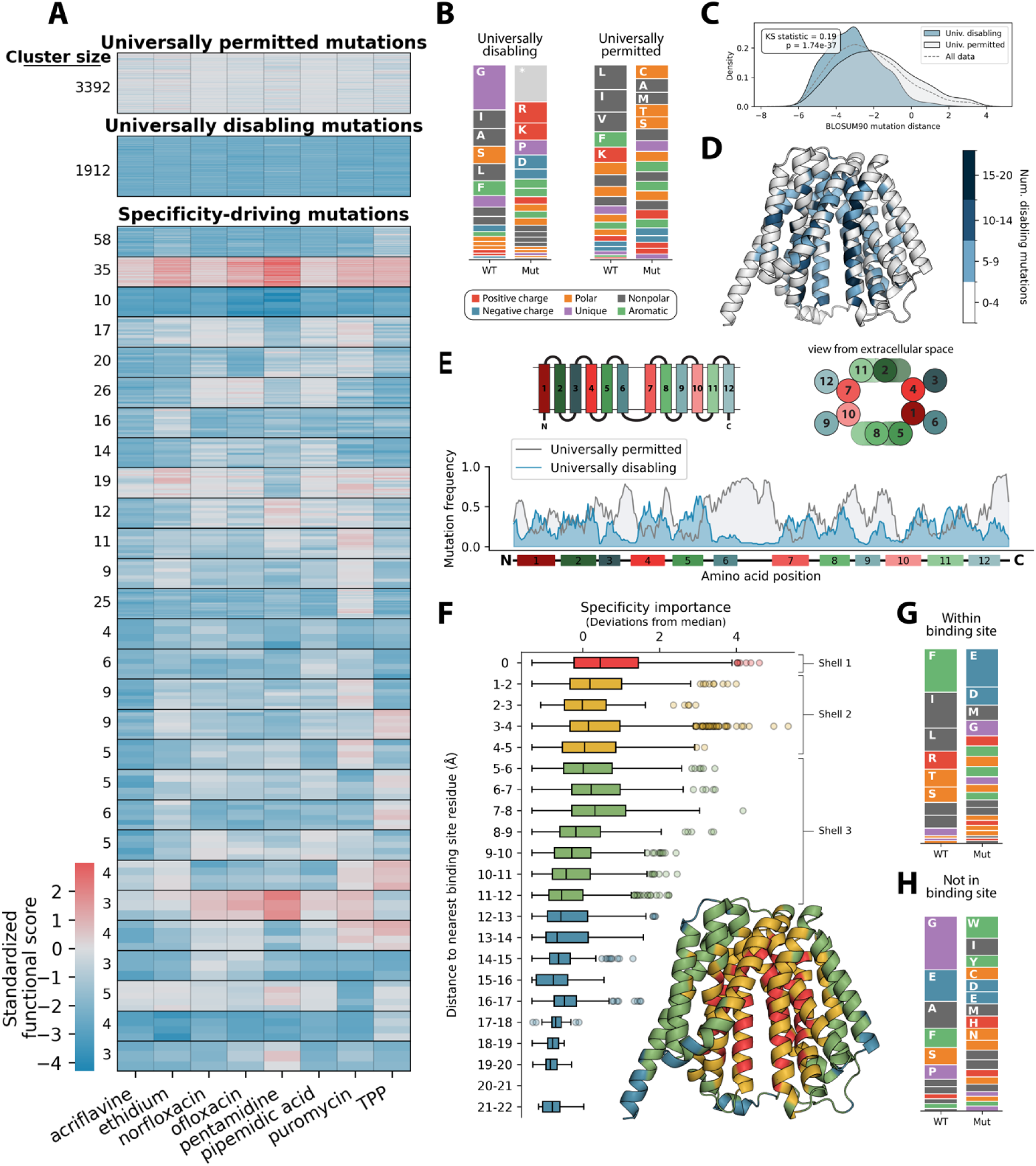
Novel insights into the molecular rules of polyspecificity. [A] Unsupervised agglomerative hierarchical clustering reveals groups of mutations with similar effects on specificity. Clusters with fewer than 3 members are omitted. Clusters that separate substrate groups with a normalized linkage distance above the first quartile are identified as specificity-driving (see Methods). [B] Amino acid frequences before and after mutation for variants in the universally permitted and universally disabling clusters. [C] Distribution of BLOSUM90 scores for mutations in the universally disabling (blue) and universally permitted (gray) clusters. [D] Number of mutations belonging to the universally disabling cluster, mapped onto the 3D structure of NorA (AlphaFold). [E] Moving average of mutation frequency across the primary sequence of NorA for both universally disabling (blue) and universally permitted (gray) mutations. [F] Specificity importance of variants grouped by distance to the nearest binding site-exposed residue. Binding site-exposed residues were determined by direct exposure to the pore in the inward-open conformation (AlphaFold). [G-H] Amino acid frequences before and after mutation for specificity-driving mutations (high outlier SI scores) within the binding site (G) or not in the binding site (H).

Before analyzing the data, we confirmed that our results recapitulate the known biology of NorA and MFS transporters. Our findings align with three known key properties: First, NorA harnesses the PMF using two essential acidic residues, Glu222 and Asp307, which bind protons regardless of the substrate being transported^34^. As expected, all non-acidic mutations to Glu222 and Asp307 were universally disabling. Second, mutations within the five MFS sequence motifs, which facilitate essential catalytic and structural features of transport^43–46^, also have a universally disabling profile. For example, Asp63 (Motif A “GxLaDrxGrk”, conformational stability) and Arg98 (Motif B “RxxqG”, proton coupling) were universally essential. Third, the disabling cluster was enriched for stop codons, charged residues, prolines, and bulky substitutions, all of which are known to disrupt membrane protein folding and stability. Mutations disrupting glycine, alanine, and serine—which are commonly found at helix-helix interfaces and are critical for compact packing of TM helices in membrane proteins^47^—were also universally disabling (Fig. 2B). In contrast, universally permitted mutations typically substitute chemically inert amino acids that are less likely to disrupt structure or function. We confirmed that variants in this cluster are less severe using BLOSUM90 distances, a measure of amino acid similarity based on evolutionary substitution patterns^48^ (Fig 2C; Kolmogorov-Smirnov statistic = 0.19, p = 1.74×10^−3^^7^).

Having established that our data align with prior knowledge, we next leveraged the DMS dataset to uncover new insights into the determinants of specificity and function. We first examined the strong positional dependence of universally permitted and disabling mutations. Disabling mutations generally occur within TM helices, especially core helices, and at bundle interfaces (Fig. 2D). Conversely, permitted mutations are common in unstructured loops and distal support helices (TMs 3, 6, 9, and 12) (Fig 2E). This agrees with evolutionary conservation, which shows higher variation in support helices compared to core helices (Supplementary Fig. 12A)^49^. These patterns suggest a modular organization of function in NorA and other MFS transporters. Peripheral helices act as malleable scaffolds that are tolerant to mutation, accommodating changes in membrane composition, while core helices facilitate catalysis and conformational change and are much more sensitive to mutation.

We next sought to identify the “functional hotspots” of NorA. Functional hotspots are positions that, when mutated, directly impair function without causing the protein to misfold. Identifying these hotspots first required us to distinguish mutations that disrupt protein stability from those that specifically affect transport activity. To determine which mutations were destabilizing, we used Rosetta to compute the ΔΔG of every variant^50^. Because NorA’s structure differs significantly across conformations, we calculated ΔΔG in three experimentally solved conformations: outward-open, occluded, and inward-open (7LO8^4^, 9B3L, and 9B3M^35^). Variants with ΔΔG exceeding one standard deviation in all three conformations were classified as misfolded. As expected, the universally disabling cluster was enriched for misfolding mutants (Supplementary Fig. 12B). However, 62.8% (974/1,550) of these mutations were predicted to remain well-folded. Thus, these mutations affect residues that are potential functional hotspots. These 974 mutations represent 200 residues; of these, we classified functional hotspots as those where five or more missense mutations abolished activity without causing misfolding (Supplementary Fig. 12C). We acknowledge that this threshold is somewhat arbitrary; the most stringent definition of a hotspot would require that nearly all substitutions eliminate function. However, few residues meet such strict criteria due to the partial activity of some mutations and accuracy of Rosetta ΔΔG predictions. This threshold balances strictness with practical sensitivity in identifying residues likely to impact function.

Alongside expected hotspots at residues involved in coupling, gating, conserved motifs, and helix-helix interfaces, we identified 16 novel functional hotspots (Supplementary Table 2). For example, Tyr278 promotes accessibility of coupling residues by forming a hydrogen bond with the Val301 backbone carbonyl, displacing TM9 and increasing solvent accessibility of Glu222 and Asp307—likely critical for proton binding (Supplementary Fig. 12D). Separately, six residues (Leu30, Asp39, Ser226, Leu227, Pro237, Asp352) at the extracellular rim may secure the inward-open conformation, mirroring the established role of cytosolic rim residues in the outward-open state^51^. Furthermore, three essential prolines (Pro56, Pro110, Pro311) create helix kinks that prevent TMs 2, 4, and 9 from obstructing the cytosolic pore^52,53^. Residues at hinge points of the N– and C-terminal bundles (Met52, Thr245, Gln255) are also essential, likely facilitating bundle rotation during conformational change^54^. Lastly, Gln51, Thr211, and Asn332 interact with NorA inhibitors in molecular dynamics (MD) simulations^55–58^, suggesting a role in a general substrate recognition mechanism and highlighting them as promising drug targets.

### Specificity-driving mutations are not restricted to the binding site

Unlike conventional protein-ligand interactions, where specificity is dictated by local binding-site residues and remains relatively narrow, we hypothesized that NorA’s broad specificity is driven by multidomain interactions that include distal regions of the protein. To test this, we computed a specificity importance (SI) score for each mutation, to identify which residues play direct roles in specificity. The SI score measures the range of functional scores across all substrates (F_max_ − F_min_), with higher scores indicating the mutation efficiently exports some substrate(s) but not others. SI scores are reported as standard deviations from the median.

We identified three “shells” of residues that govern specificity (Fig. 2F). The first shell (red) directly interacts with substrates, playing an expectedly major role in specificity through direct molecular recognition (90th percentile SI = 2.6, maximum = 4.6). The second shell (yellow) contains residues 1–5Å from the binding site (90th percentile SI = 1.7, maximum = 5.3). These likely contribute to specificity by modulating energy coupling—for example, G251D, which adds a negative charge near coupling residues and is tolerated by ethidium only (Supplementary Fig. 13A)—or by impacting binding site flexibility, as in the puromycin-specific mutation G147W, which disrupts a helix-helix interface 4.4Å from the binding site (Supplementary Fig. 13B).

Surprisingly, the third shell (green, 5–12Å from the binding pocket) still influences specificity, despite its distance (90^th^ percentile SI = 1.3, maximum = 4.2). These residues may affect specificity through several mechanisms: (i) allosteric regulation, where mutations alter binding pocket dynamics via long-range conformational coupling; (ii) structural stabilization, where residues contribute to overall fold and relative stability of conformational states, indirectly shaping substrate interactions^59^; and (iii) membrane interactions, where lipid contacts influence transporter flexibility and substrate access^60,61^. For example, E356P (SI = 2.2, 11.2Å from the binding site) may dictate the lipid embedding of TM12 by removing the negative charge at the extracellular terminus of this TM helix^62^ (Supplementary Fig. 13C), while I177W (SI = 3.3, 8.9Å from the binding pocket) may obstruct interactions between N– and C-terminal bundles that stabilize the outward-open state^63^ (Supplementary Fig. 13D). These findings suggest NorA’s broad specificity is not solely encoded by the binding site but emerges from a complex interplay of residues across multiple structural layers.

We hypothesized that distal mutations regulate specificity by fundamentally different mechanisms than those occurring within the binding site. To test this, we compared specificity driving mutations (high outlier SI scores) by their binding-site exposure. Binding-site mutations primarily alter hydrophobicity (e.g., isoleucine substitutions from TM1) or substitute residues that directly interact with substrates (Phe303 and Phe306)^4,64^. Arg310, which may be involved in proton translocation, also showed substrate-dependent importance, suggesting a link between energetics and specificity. This is further supported by frequent addition of acidic residues, which may affect coupling efficiency^19^ (Fig. 2G). In contrast, distal mutations often involve glycine and alanine substitutions at bundle interfaces, likely disrupting structural or conformational features. Also common are mutations at Phe85, which modulates pK_a_ of Arg98^34,37^ (coupling), and Lys67, which forms a charge-relay triad with Asp63 and Asp118^63^ (conformational stability), underscoring the role of diffuse networks of interacting residues in polyspecificity (Fig. 2H).

Together, these analyses reveal that substrate specificity in NorA emerges from a highly distributed network of residues spanning the binding site, adjacent helices, and distal structural elements. Specificity-driving mutations occur not only at direct contact points but also at remote sites that influence coupling, conformational dynamics, and membrane interactions. This modular organization underlies NorA’s extraordinary functional adaptability.

### Novel insights into the mechanistic basis of polyspecificity

To better understand the structural principles underlying substrate specificity, we examined how sequence changes drive distinct specificity profiles. We highlight six examples that illustrate how mutation can reshape substrate selectivity.

One cluster of mutations conferred resistance to puromycin only (N = 25, Fig. 3B). These mutations are located at hydrophobic residues near NorA’s proton-binding sites, particularly at Ile244 (Supplementary Fig. 13E). In transporters, this hydrophobic environment is critical for tuning the pK_a_ of proton-binding residues into physiological range^65^. Puromycin, which has a relatively high topological polar surface area (TPSA) compared to other substrates (TPSA = 161Å^2^, other substrates average = 77.1Å^2^), may favor a less hydrophobic binding pocket. Because proton binding and substrate binding are often negatively linked^66^, stronger interactions with puromycin could facilitate proton release, enabling efflux even when the local hydrophobicity is disrupted.

**Figure 3.**
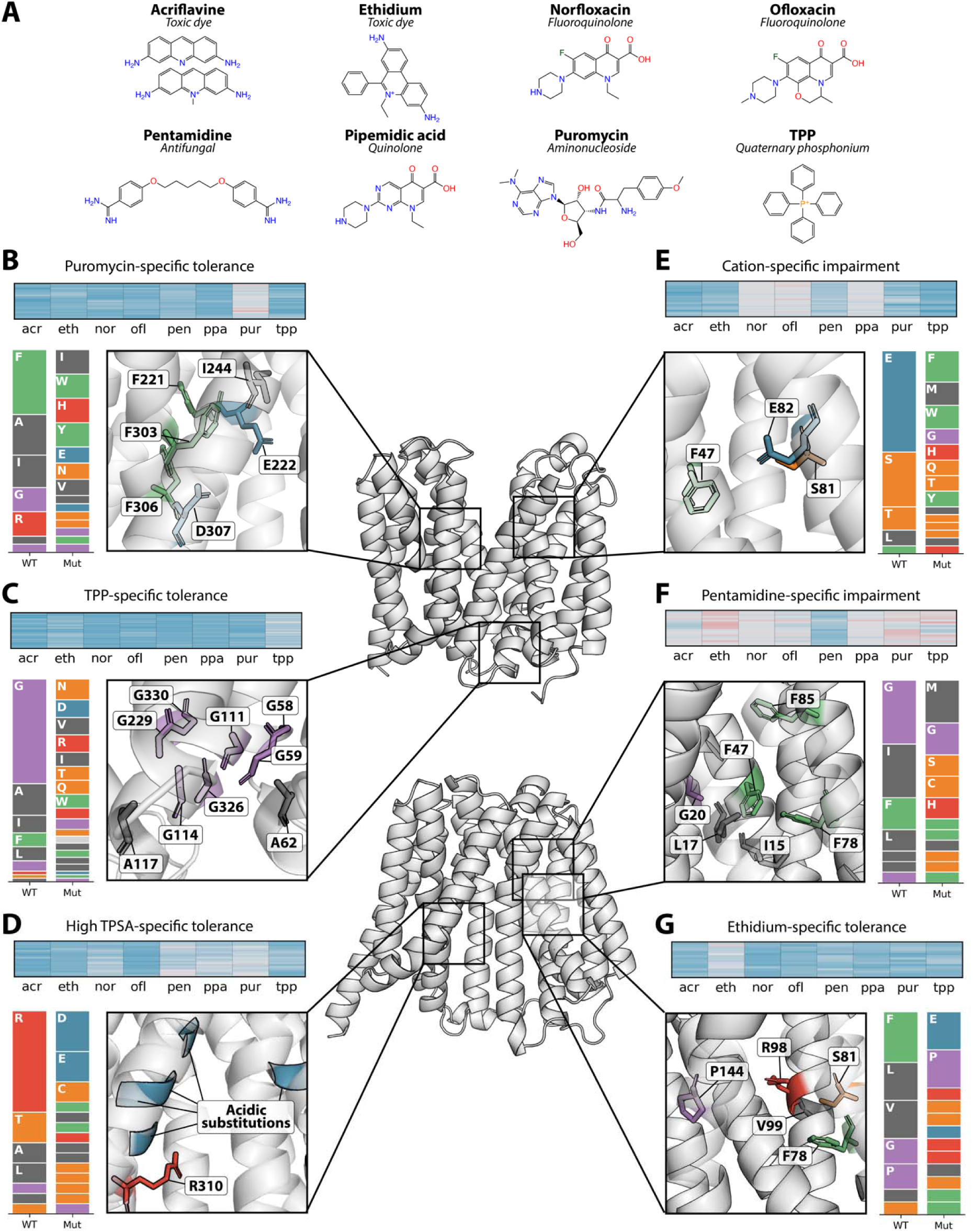
Structural mechanisms of select specificity-driving clusters. [A] Chemical structures for each of the eight substrates used to select the NorA DMS library in this work. [B-G] Local heatmaps for groups of mutations identified by unsupervised hierarchical clustering, distributions of wild-type and mutant amino acids, and select positions on the NorA structure (7LO8 and AlphaFold) to illustrate the potential mechanism. Abbreviations: acr (acriflavine); eth (ethidium); nor (norfloxacin); ofl (ofloxacin); pen (pentamidine); ppa (pipemidic acid); pur (puromycin); tpp (tetraphenylphosphonium).

By contrast, many variants conferred resistance to TPP alone (N = 58, Fig. 3C). These mutations predominantly involved bulky substitutions at glycine and alanine residues, particularly at bundle interfaces (Supplementary Fig. 13F). At the cytosolic face, these small amino acids normally allow close association of N– and C-terminal bundles, enabling Asp63 to form a hydrogen bond with the backbone amide of Arg324, which stabilizes the outward-open conformation^63,66^. TPP, one of the smallest and most lipophilic substrates we tested (logP = 6.63, other substrates average = 0.24), may not require a fully open pore for transport. Instead, its high lipophilicity could allow it to partition into the membrane and access the binding site through a more compact or partially occluded conformation^67,68^.

In a distinct pattern, a third group of variants retained the ability to transport only pentamidine, pipemidic acid, and puromycin (N = 20, Fig. 3D). Most of these mutations disrupt Arg310, either through direct substitution or charge neutralization by nearby acidic mutations (Supplementary Fig. 13G). Arg310, a basic residue in the binding site, is implicated in promoting proton release in homologous transporters^37^ and directly interacts with Asp307 in the inward-open conformation (9B3M)^35^. Pentamidine, pipemidic acid, and puromycin have higher polar surface area (TPSA = 118, 98.7, and 161Å^2^) compared to other substrates (TPSA average = 51.6Å^2^). This polarity may partially compensate for the loss of Arg310, facilitating proton transfer through alternative interactions.

Variants in another group (N = 26, Fig. 3E) failed to transport cationic substrates. Most mutations substituted Glu82 or adjacent residues, suggesting that this buried negative charge plays a role in facilitating cation transport^69^ (Supplementary Fig. 13H). Supporting this idea, we observed that the conservative substitution E82D was tolerated, and that F83D enhanced activity specifically toward cationic substrates. These findings are consistent with a model in which a localized negative charge promotes transport of cationic substrates, potentially through indirect charge-charge interactions.

In addition, another set of variants (N = 19, Fig. 3F) retained transport of all substrates except pentamidine. These mutations generally involved modest disruptions to structural features in core helices, such as replacement of interfacial glycines to short-chain amino acids or nonpolar to polar substitutions (Supplementary Fig. 13I). Such changes likely produce a smaller or more polar binding pocket. Given pentamidine’s large size and hydrophobicity, it may be preferentially excluded from these altered environments^70^.

Finally, a distinct set of variants (N = 16, Fig. 3G) shifted specificity exclusively to ethidium. These mutations frequently introduced a charged residue, particularly near the proton-coupling residue Arg98. Mutations to Pro144, part of the conserved motif C (“GPxxGG”), are also found in this cluster (Supplementary Fig. 13J). Motif C is thought to link proton transfer to drug antiport while preventing opportunistic symport^71^. Charged mutations near Arg98 may attenuate its electrostatic role in coupling. These alterations may impair coupling or conformational dynamics in a way that selectively permits ethidium export.

These examples collectively highlight the remarkable adaptability of NorA: subtle sequence changes at different sites can reshape substrate specificity in highly specific and predictable ways. Although multidrug transporters are often described as promiscuous, the molecular basis for their substrate selectivity has remained poorly understood. Our DMS approach systematically uncovers the structural and biochemical logic underlying these changes, revealing how local perturbations—whether in charge, hydrophobicity, or conformational flexibility—can bias transport toward distinct classes of compounds.

### Sensitivity to basic extracellular pH as a proxy for energetic efficiency

The relationship between substrate specificity and transport efficiency is a fundamental but underexplored question in transporter biology. Transporters consume energy to perform the work of substrate efflux, and promiscuous transporters must accommodate the energy demands of many potential substrates. Previous work has shown that energy coupling is flexible and may vary across substrates, but the energetic costs of broad versus narrow specificity remain largely unexplored^18–25^.

Similar principles have been observed in other molecular machines, like cytochrome P450 enzymes balancing catalytic efficiency with substrate diversity^72^ or ribosomes tuning translation speed and fidelity based on energetic constraints^73^. Drawing on these parallels, we sought to define the core principles linking specificity and efficiency in transporters.

Two competing hypotheses address the relationship between transporter specificity and efficiency. The first argues that promiscuous transporters are inherently inefficient, because the flexibility required to bind many substrates compromises the conformational precision needed for efficient energy coupling. This is observed in ATP-driven efflux pumps, where promiscuous transporters have higher basal ATP hydrolysis than more specific homologs^28^. The alternative hypothesis proposes the opposite: that broad-specificity transporters must be efficient. Here, high energy efficiency allows efflux of diverse substrates without needing finely-tuned binding and release energies for each—an impractical requirement for transporters that act on many compounds. Instead of evolving precisely tuned mechanisms for every substrate, efficient transporters may rely on robust energy coupling to offset variability in substrate binding affinity.

A key challenge in testing this relationship is designing an assay that can quantify transport efficiency in a pooled high-throughput format. NorA is a proton-coupled transporter driven primarily by ΔpH, the chemical component of the PMF^30,74^: as protons flow down their concentration gradient, they power substrate export against its gradient. Because intracellular pH is tightly regulated, buffering extracellular pH modulates ΔpH^75–79^. This chemical perturbation effectively limits (higher pH_out_, reduced ΔpH) or bolsters (lower pH_out_, larger ΔpH) the driving force behind NorA-mediated drug export. We hypothesized that efficient transporter variants could confer resistance even with limited ΔpH, while less efficient variants would strictly require a strong driving force. To test this, we performed selections with varied extracellular pH. Variants that confer resistance at high pH_out_ are inferred to be more efficient, while those that function only at low pH_out_ are less so, thus indirectly measuring efficiency in a pooled screen.

To map the determinants of energy efficiency and ultimately test the relationship between promiscuity and efficiency, we challenged the NorA DMS library with norfloxacin in buffered growth medium at pH 6 or pH 7. The difference in function between these two pH conditions, ΔF_pH_ (F_pH 7_ − F_pH 6_), represents the relative dependence of efflux on a strong ΔpH driving force. A negative ΔF_pH_ reflects pH sensitivity and less efficient transport than wild type (Fig. 4A, purple lines), while a positive ΔF_pH_ indicates pH tolerance and more efficient transport (Fig. 4A, orange lines).

**Figure 4.**
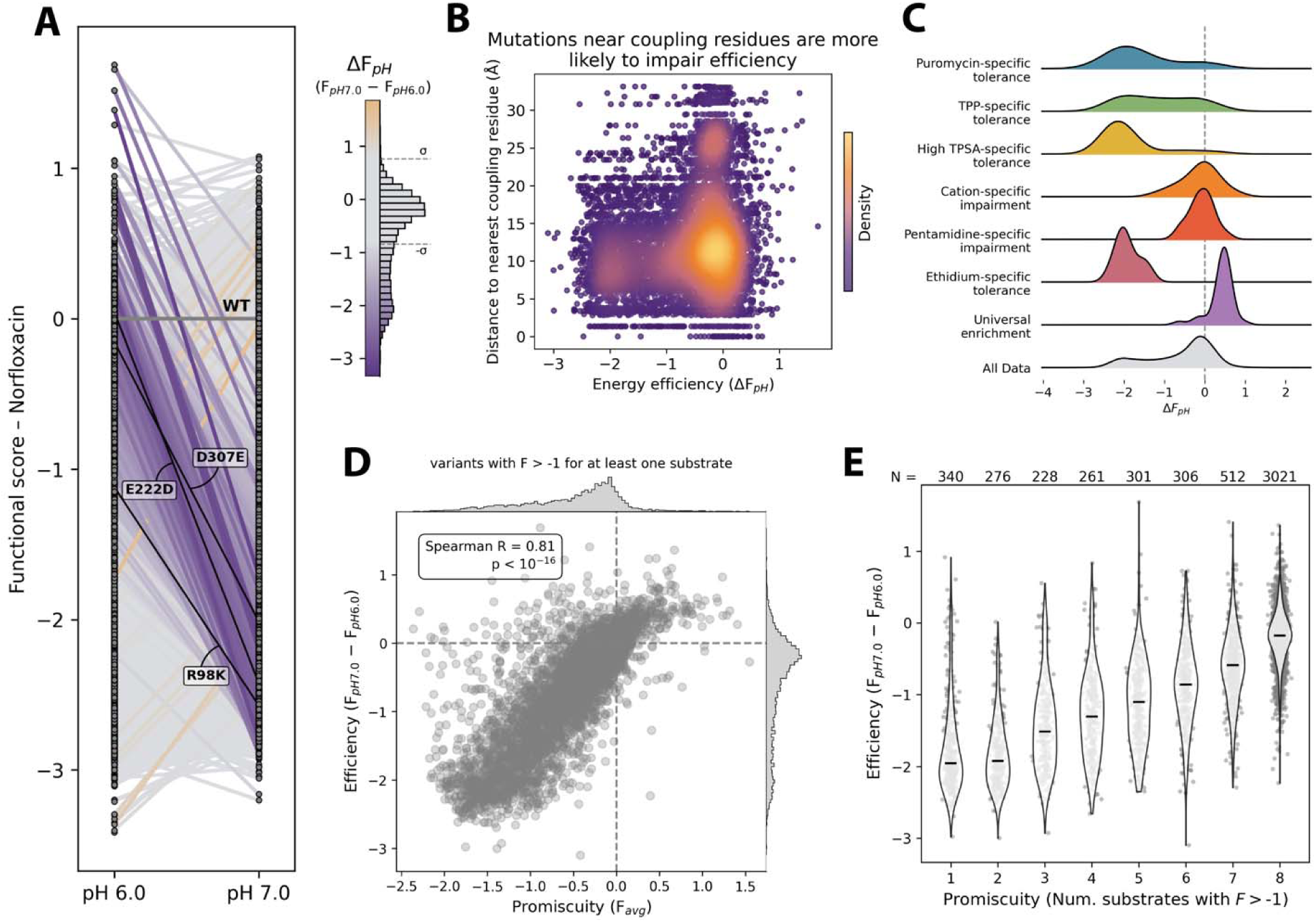
pH sensitivity of transport correlates strongly with promiscuity. [A] Norfloxacin transport activity of each variant in the NorA DMS library (gray dots) at pH 6.0 (left) and pH 7.0 (right). Lines indicate the change in each variant’s performance between pH conditions. Wild type is shown in dark gray. Variants within one standard deviation of wild type are shown in light gray, increased pH sensitivity in purple, and decreased pH sensitivity in orange. Select variants are labeled and highlighted in black. [B] Scatter plot comparing ΔF_pH_ scores with distance from the nearest coupling residue (E222, D307, or R98). [C] Distributions of ΔF_pH_ scores for each of the specificity clusters highlighted in Figure 3 (top six distributions), universally enriched variants (purple) and all data (gray). [D] Scatter plot comparing promiscuity (average functional score in all specificity screens) with efficiency (ΔF_pH_ from norfloxacin pH-sensitivity screen) for variants with activity on at least one substrate. [E] Violin plot showing typical ΔF_pH_ values for variants with activity on 1–8 substrates tested.

We first confirmed that our approach recapitulates prior knowledge of NorA energy coupling. Our results are consistent with two key expectations: First, direct mutation of coupling residues causes high pH sensitivity. Glu222, Asp307, and Arg98 play essential roles in proton binding and release. Mutations to these positions which do not fundamentally change their ability to interact with protons—E222D, D307E, and R98K—show significant energetic impairment (Fig. 4A, labeled black lines). Similarly, mutation of Asn137 to other polar residues results in strong negative ΔF_pH_ scores, consistent with its role in linking Asp307 protonation to conformational change^34^. Mutation of Arg310, a putative coupling residue, is also consistently pH sensitive (Supplementary Fig. 14A). Second, mutations proximal to coupling residues are more likely to impair efficiency. We computed each mutation’s distance to the nearest proton coupling residue (Glu222, Asp307, or Arg98) and compared this distance to ΔF_pH_ scores. As expected, mutations near coupling residues are more likely to impair efficiency, while distal mutations very rarely cause this phenotype (Fig. 4B).

After confirming that high-throughput pH sensitivity aligns with prior knowledge of proton coupling, we probed the data for novel understanding of energy efficiency in NorA. While the function of most variants did not change between pH 6 and pH 7 relative to wild type, 16.8% (N = 1,306 variants) had significant energetic impairment (Supplementary Fig. 14B; FDR-adjusted p < 0.05, ΔF_pH_ < –2σ). Many of these mutations disrupt the environment of coupling residues by adding proximal charges or altering charge buriedness (e.g., Gly101 mutations). Interestingly, several efficiency-impaired mutants are found at the cytosolic loop connecting N– and C-terminal bundles (Glu192, Pro193, and Gln194). Some variants with gain-of-function efficiency phenotypes also appear here (K181T, S183N, T185V, and F188C), supporting a role of the amphipathic helix within this loop in sensing protonation^63,80^. We also observed energetic defects for 11-, 12– and 13-amino acid C-terminal truncation mutants (E376*, K377*, and Q378*), similar to the Δ107 truncation mutant of EmrE (a model SMR-family transporter) which exhibits a severe coupling defect^81^.

We next examined the efficiency measurements in the context of altered specificity (see Fig. 3). We compared the ΔF_pH_ values of mutations found in each specificity cluster (Fig. 4C). Mutations associated with puromycin-specific tolerance (blue, pK_a_-tuning of proton binders), high TPSA-specific tolerance (yellow, Arg310 neutralization), and ethidium-specific tolerance (dark red, Arg98 attenuation) have strongly negative ΔF_pH_ scores, consistent with the proposed energetic basis of altered specificity for these clusters. In contrast, ΔF_pH_ scores remained neutral for clusters with mechanisms not related to coupling (cation-specific impairment: orange, electrostatic attraction; pentamidine-specific impairment: red, binding pocket size and polarity). Multiple mechanisms likely contribute to the TPP-specific tolerance cluster (green), as both negative and neutral ΔF_pH_ scores are seen here. Strikingly, mutations that showed universal enrichment (purple) have positive ΔF_pH_ scores, suggesting that enhanced energy efficiency underlies substrate-independent gain-of-function.

### Efficient energy coupling enables promiscuity in NorA

We next compared efficiency with the specificity breadth of each variant, leveraging our dataset from the eight-substrate screen. This comparative analysis directly tests the two competing hypotheses: whether broad specificity comes at an energetic cost or if high efficiency is a prerequisite for polyspecificity.

We calculated the average functional score across all eight substrates (F_avg_) as a measure of promiscuity and compared ΔF_pH_ to this value. To definitively exclude misfolded variants, which are inactive regardless of pH, we limited this analysis to mutants with activity on at least one substrate (N = 5,251). We observed a strong positive correlation between efficiency and promiscuity (Fig. 4D; Spearman R = 0.81, p < 10^-16^). Mutations that narrow specificity are less efficient (bottom left), while those with activity on all substrates are more efficient (top right).

Although some variants are both efficient and specific (top left), inefficient yet promiscuous variants are extremely rare (bottom right), supporting a model where energy efficiency is necessary but not sufficient to achieve broad specificity.

As an additional intuitive measure of promiscuity, we calculated the number of substrates for which each variant exhibited significant activity (F > –1) and compared this value to ΔF_pH_ (Fig. 4E). Again, variants with activity on more substrates have higher ΔF_pH_ scores (higher energy efficiency) than more specific variants. Efficiency as measured by pH sensitivity of acriflavine transport shows similar results (Supplementary Fig. 14C, D).

### Clonal measurements validate high-throughput pH sensitivity

To validate the efficiency measurements from our pooled screen, we performed clonal dose-response assays with norfloxacin in buffered growth media. IC_50_ values calculated from these assays reflect the magnitude of efflux-mediated resistance, mirroring functional scores F_pH 6_ and F_pH 7_. The difference in resistance across pH directly measures each variant’s dependence on ΔpH, grounding the pooled screen in traditional microbiological methods.

We built 14 NorA variants individually and collected norfloxacin dose-response curves for MG1655-ΔacrAB *E. coli* expressing each variant in isolation at pH 6 or pH 7. IC_50_ values calculated from these curves agreed with our high-throughput assay: relatively pH-tolerant variants like wild-type NorA (ΔF_pH_ = 0) showed a smaller shift in IC_50_ between pH 6 and pH 7 than pH-sensitive variants like A105E (ΔF_pH_ = –2.4) (Fig. 5A). For each of the 14 variants tested clonally, we calculated the variant’s effect relative to wild type (F_clonal_ = log_2_(IC_50, var_ / IC_50, WT_)), then computed a clonal ΔF_pH_ (ΔF_pH, clonal_ = F_clonal, pH 7_ − F_clonal, pH 6_). The IC_50_-based ΔF_pH, clonal_ scores correlated well with the high-throughput sequencing-based ΔF_pH_ values (Spearman R = 0.77, p = 0.001), validating our pooled screen’s ability to report pH sensitivity of NorA-mediated drug resistance (Fig. 5B).

**Figure 5.**
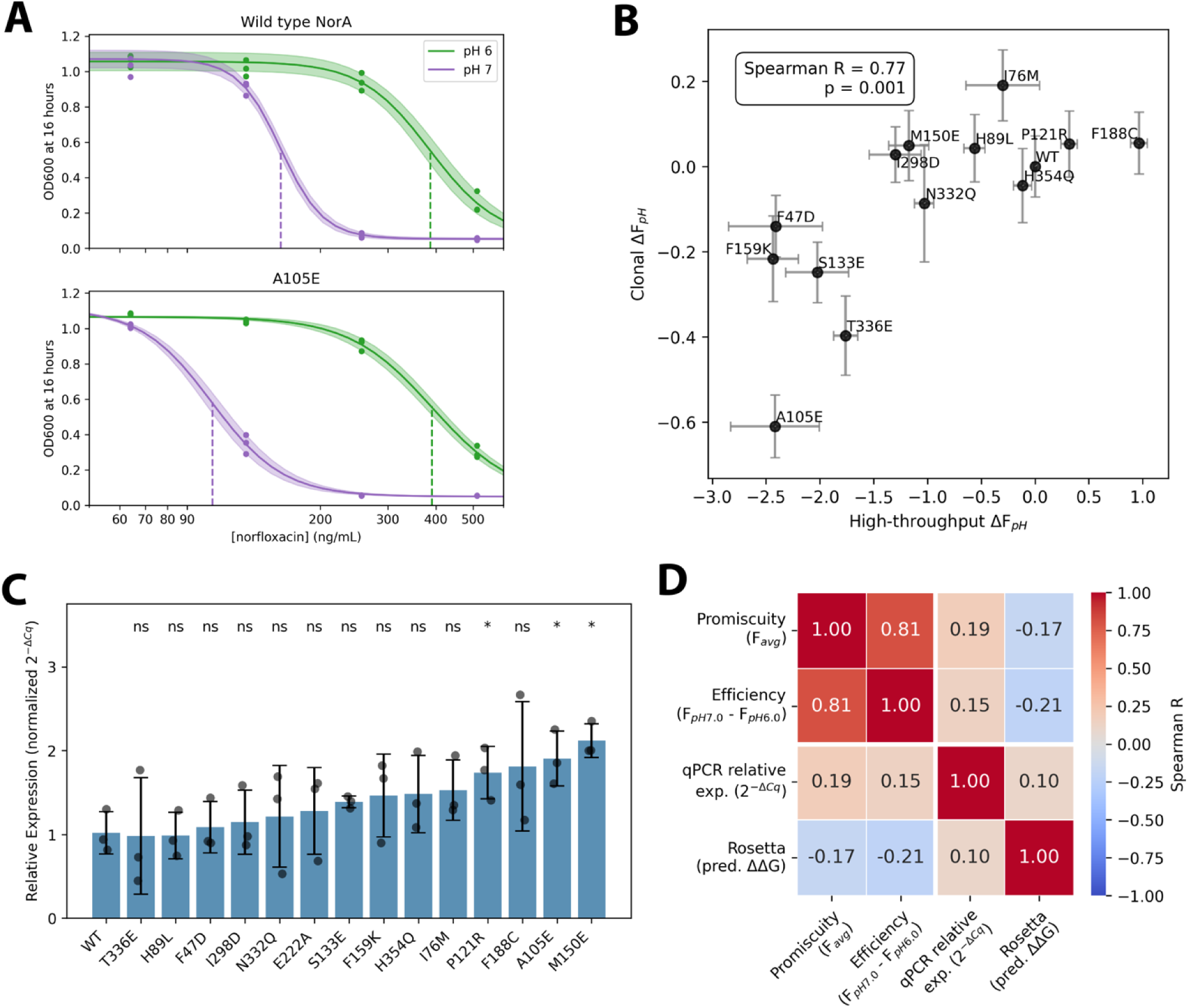
Clonal validation of high-throughput pH sensitivity measurements and RNA– and protein-level abundance. [A] Representative pH-tolerant variant (wild type) and pH-sensitive variant (A105E) IC_50_ curves. IC_50_ values are marked with dotted lines. [B] Correlation of high-throughput ΔF_pH_ with clonally measured IC_50_-based ΔF_pH_ validates the high-throughput pH-sensitivity assay. [C] Clonally measured RT-qPCR relative expression levels show that significant RNA-level abundance differences are uncommon and modest. [D] Neither RNA-level abundance differences (qPCR relative expression) nor protein-level stability differences (Rosetta predicted ΔΔG) correlate significantly with promiscuity or efficiency.

Because mutation can alter mRNA or protein abundance, an alternative explanation for our findings is that differences in variant abundance cause changes in drug resistance, rather than specific functional effects. To evaluate the role of RNA-level abundance in our measurements, we determined transcription differences for the 14 clonally constructed variants and wild type using reverse transcriptase quantitative PCR (RT-qPCR). Only 3 mutants had altered transcription relative to wild type, with at most a 2.1-fold increase in expression level (Fig. 5C). These modest differences did not correlate with promiscuity or pH sensitivity. In consideration of protein-level stability differences, we compared our Rosetta ΔΔG calculations with functional measurements. Again, these values did not correlate with promiscuity or pH sensitivity (Fig. 5D). We conclude that, while some abundance and stability differences do exist within the library, they cannot explain the functional differences and relationships uncovered here.

### Thermodynamic model of efflux pump specificity

Our mutational screen of NorA revealed that variants with high energy efficiency also exhibit the broadest substrate specificity. To explain this, we propose a thermodynamic model of proton-coupled transport. Efflux pumps like NorA harness the PMF to power substrate efflux against a concentration gradient^63^. The total free energy available from the PMF is given by:

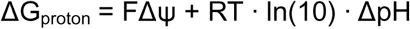

where Δψ is the membrane potential (inside relative to outside), ΔpH = pH_out_ − pH_in_, F is the Faraday constant, R is the gas constant, and T is temperature. This energy can be partially converted into productive work—i.e., translocation of substrate—by the transporter. The effective free energy available for transport is thus:

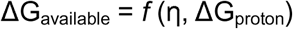

where η is the coupling efficiency, an intrinsic property of the transporter that reflects how effectively it transduces the PMF into mechanical work (Fig. 6A). η depends on factors such as the pK_a_ of proton-binding residues and the likelihood of conformational change in the proton-bound state.

**Figure 6.**
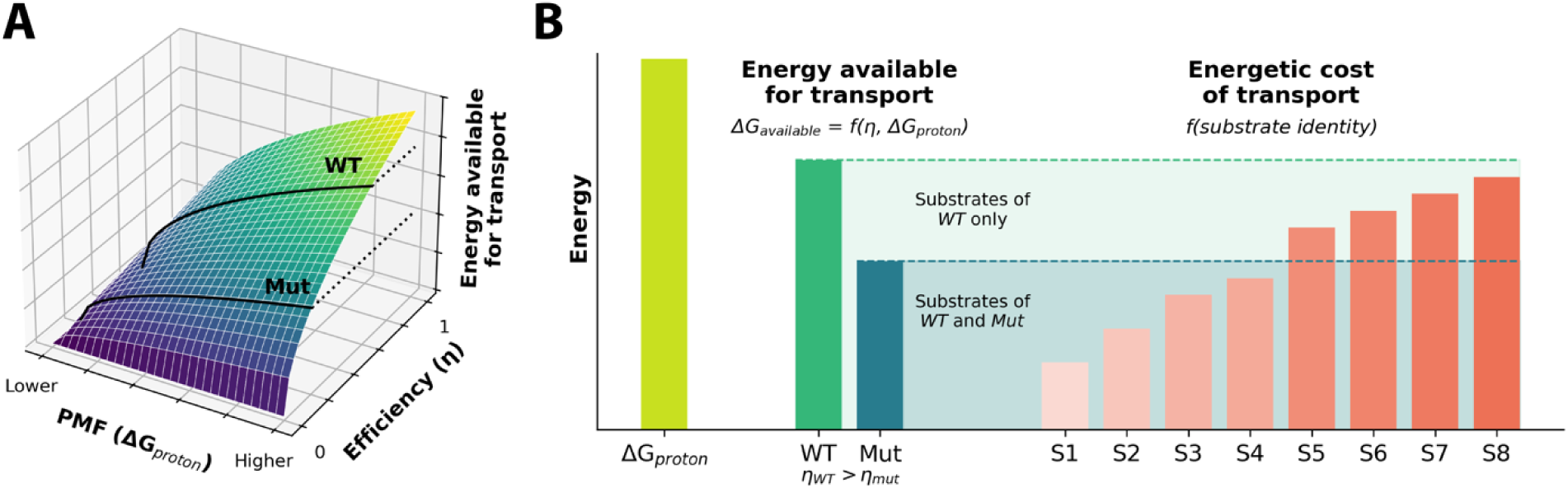
A theoretical thermodynamic model for the relationship between promiscuity and energy efficiency. [A] Energy available for transport (ΔG_available_, z-axis) is a function of the strength of the driving force (ΔG_proton_) and the ability of the transporter protein to extract that energy (η). [B] Because different substrates will have different energetic costs (red shaded bars), inefficient variants with less energy available for transport (blue bar) are only able to confer resistance to the “cheaper” substrates and are necessarily less promiscuous than more efficient variants with excess energy available for transport (green bar).

Each substrate has its own transport cost, ΔG_substrate_, determined by binding enthalpy and translocation energy. Transport is thermodynamically feasible only if:

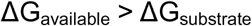

To focus on variation attributable to the protein sequence, we assume a constant effect of Δψ across all variants. Under this assumption, sequence-dependent differences in η dominate variation in transport efficiency. High-η variants have a larger energetic “budget,” allowing them to efflux a broader range of substrates, including those with high transport costs. Conversely, low-η variants are limited to substrates that are energetically cheap to transport (Fig. 6B).

Thus, efficiency is a prerequisite for promiscuity: only high-efficiency transporters can overcome the energetic barriers posed by a diverse set of substrates. Our model explains why the most energy-efficient NorA variants in our screen also exhibit the greatest functional promiscuity.

## Discussion

Our findings reveal that the molecular determinants of transporter specificity extend beyond the binding site to a distributed network of residues involved in energy coupling, membrane interactions, and conformational dynamics. We emphasize that substrate binding alone does not guarantee productive transport: efficient efflux requires coordinated control of protein and environmental energetics, the rate and magnitude of conformational transitions, and dynamic binding and release of substrate and protons. The specific conditions required for each of these factors vary depending on substrate size, antimicrobial potency, or structural and chemical features. To achieve polyspecificity, efflux pumps operate above the minimal energetic and structural thresholds required for individual substrates. As a result, mutations can drive specificity by altering energy efficiency, conformational flexibility, or coupling kinetics, even without directly disrupting substrate binding. Together, these findings uncover general principles by which multidrug transporters balance broad molecular recognition with efficient energy transduction.

To comprehensively map the molecular determinants of specificity, we performed deep mutational scanning of the NorA efflux pump with eight substrates and two proton gradient conditions. NorA, which is commonly overexpressed in MDR infections like MRSA, confers resistance to numerous clinically relevant compounds, making it a critical model for understanding general principles of drug efflux. Our screen identified 1,912 universally disabling mutations, which impair protein folding or disrupt known and novel functional hotspots, and 3,392 universally permitted mutations, largely within flexible loop regions. We also uncovered numerous specificity-driving mutations, both proximal and distal to the binding pocket, that selectively reprogram transport activity toward distinct substrates. Unsupervised clustering revealed an eight-way specificity switch, demonstrating that different substrates impose independent structural and energetic requirements. By measuring pH-dependent transport efficiency, we further uncovered a fundamental relationship between energy efficiency and promiscuity: efficient variants have broad specificity profiles, whereas inefficient variants are restricted to narrow substrate ranges. Together, these data establish a direct link between energy efficiency and substrate promiscuity, defining key principles underlying polyspecific efflux.

High-throughput studies of drug efflux pumps are beginning to enable a scale of functional information previously unattainable^82^. However, we acknowledge a fundamental limitation of these approaches: to enable selection-based screening, function must be linked to a selectable phenotype such as viability, inherently compressing mechanistic nuance into a single readout. To address this, we used multiparametric assays that deconvolute growth effects into distinct elements of transporter function, including substrate efflux (drug resistance), energy efficiency (pH sensitivity), and protein stability (Rosetta ΔΔG). Nevertheless, key aspects of transporter activity remain challenging to resolve at scale. In particular, methods to systematically measure conformational rates and kinetic coupling in high throughput have not yet been developed. Extending high-throughput functional screens to directly capture transporter dynamics represents a major opportunity for the field and would provide a powerful complement to static structural and functional analyses.

The data presented here lay a foundation for advancements in biotechnology and biomedicine. Our DMS across substrates begins mapping the relationship between features of NorA’s protein sequence and its substrate range, paving the way for engineered transporters with tailored specificity. Designer transporters could optimize biosynthetic flux and resistance to reactant impurities, revolutionizing microbial synthesis of biofuels and bioproducts^83–85^. The basic knowledge built here also stands to benefit the fight against multidrug resistance in the clinic. By fully mapping NorA’s functional hotspots, we highlight drug targets to facilitate rational design of efflux pump inhibitors^86^. Additionally, our findings on energy efficiency and specificity breadth support targeting of coupling pathways to counter drug efflux, an approach that has already seen some promising results^87,88^. This study not only advances fundamental understanding of NorA’s function but also establishes a platform for engineering transporter specificity and for targeting efflux in antibiotic resistance.

**Supplementary Figure 1.**
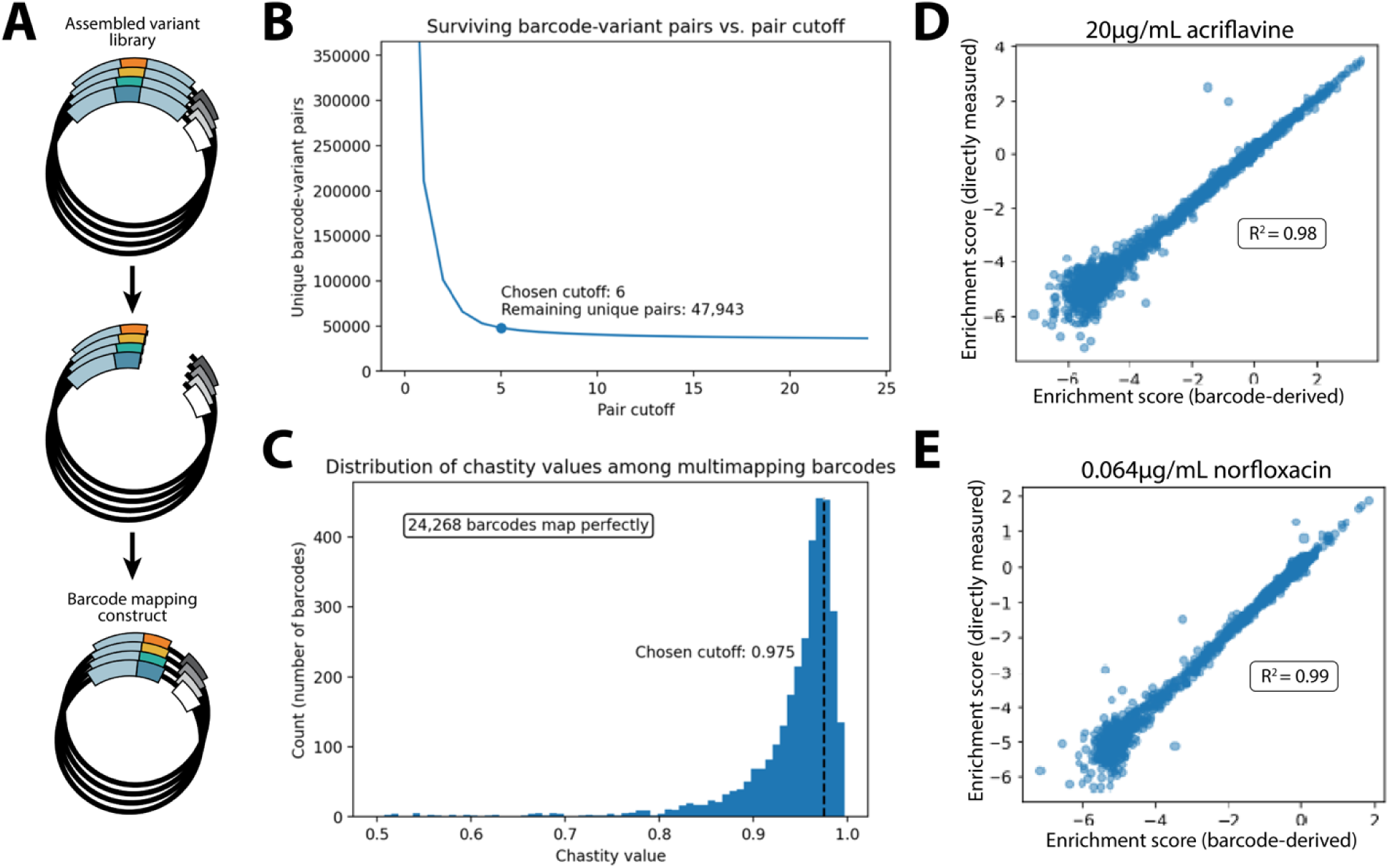
Barcoding the DMS library. [A] Construction of mapping constructs for mapping internal subpools. The variable region of each library is positioned subpool directly adjacent to barcodes using PCR and Golden Gate assembly (see Methods), allowing for short-read Illumina sequencing to associate barcodes with NorA variants. [B] Method of setting read-count cutoffs for a representative subpool. Cutoffs were set at the elbow of the plot of unique barcode-variant pairs remaining and read-count cutoff. [C] Method of removing disloyal barcodes from mapped sequences. Chastity values are calculated to quantify barcode loyalty (see Methods). Barcodes with chastity values lower than 0.975 are removed. [D-E] Empirical evidence of successful barcode mapping. Barcode-derived enrichment scores correlate strongly with enrichment scores from direct variant sequencing after selecting a single subpool with 20μg/mL acriflavine (R^2^ = 0.98) or 0.064μg/mL norfloxacin (R^2^ = 0.99).

**Supplementary Figure 2.**
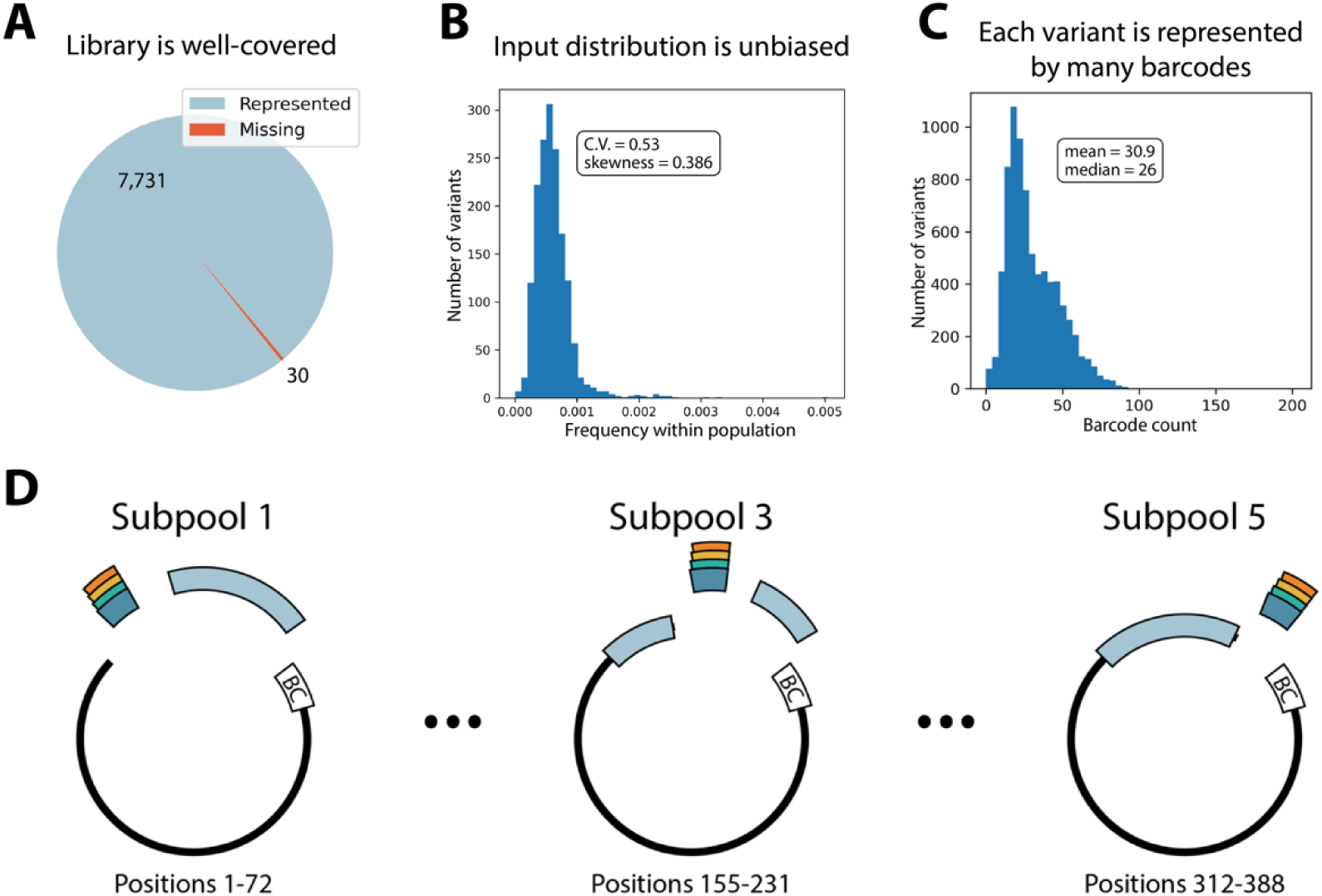
Library construction and quality. [A] 7,731 of the 7,761 expected variants were present in the library (99.7%). [B] Histogram of variant frequencies in the pre-selection library (Coefficient of variation [σ/μ] = 0.53, Fisher-Pearson coefficient of skewness = 0.386). [C] Histogram of barcode multiplicities in the NorA DMS library (mean = 30.9 barcodes per variant, median = 26 barcodes per variant). [D] NorA DMS library assembly scheme. Subpools 1-4 used three-part Golden Gate assemblies while subpool 5 only required a two-part assembly.

**Supplementary Figure 3.**
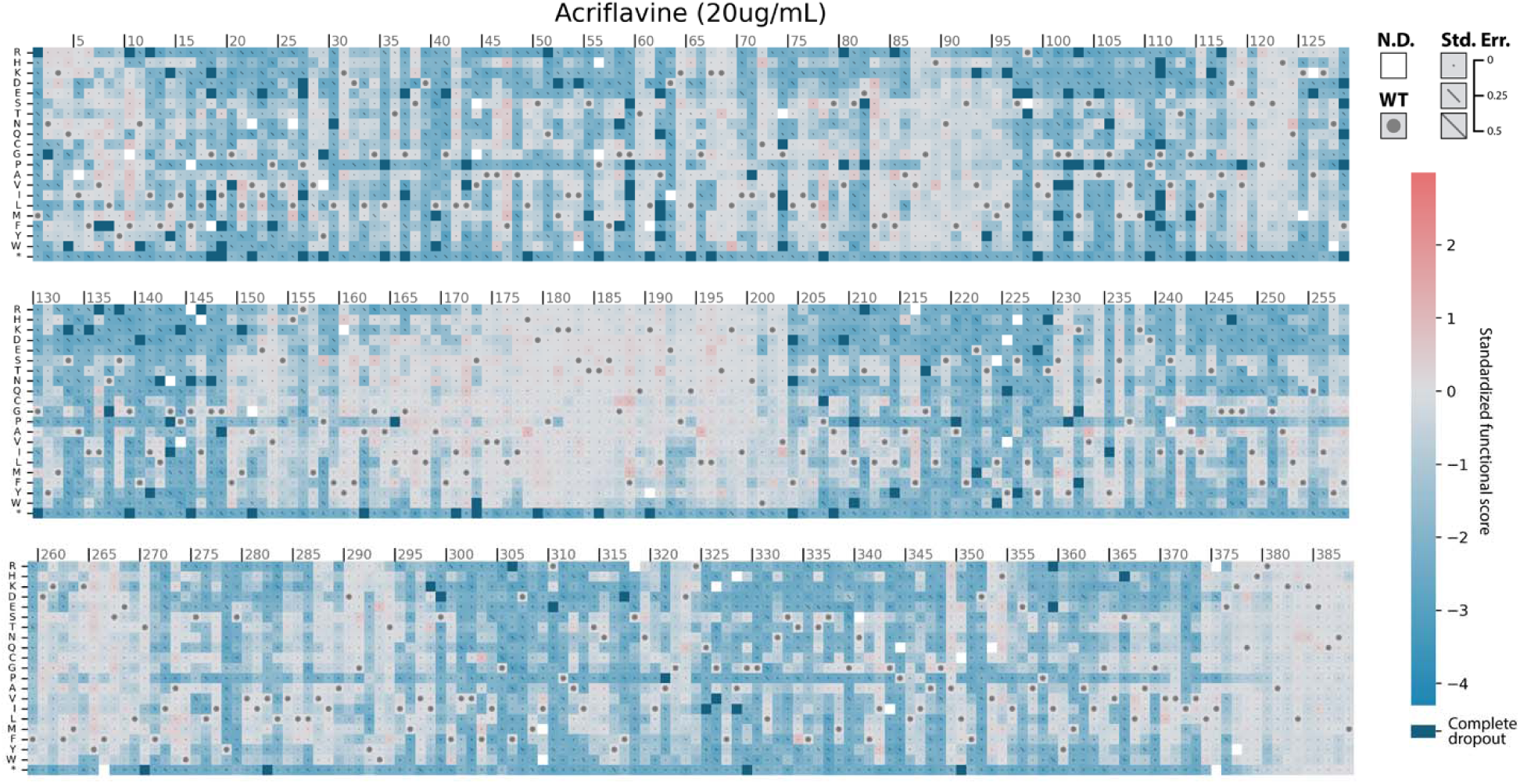
Sequence-function landscape of acriflavine transport. Functional scores for all variants after 12 hours of competitive growth in the presence of 20μg/mL acriflavine. The wild-type residue at each position is marked with a dot. Error values are shown as diagonal lines across each square, scaled so that the entire diagonal represents an error of 0.5 standardized functional score units. Variants with error > 0.5 are omitted (white). Variants which dropped out completely (zero observations after selection) are shown in dark blue.

**Supplementary Figure 4.**
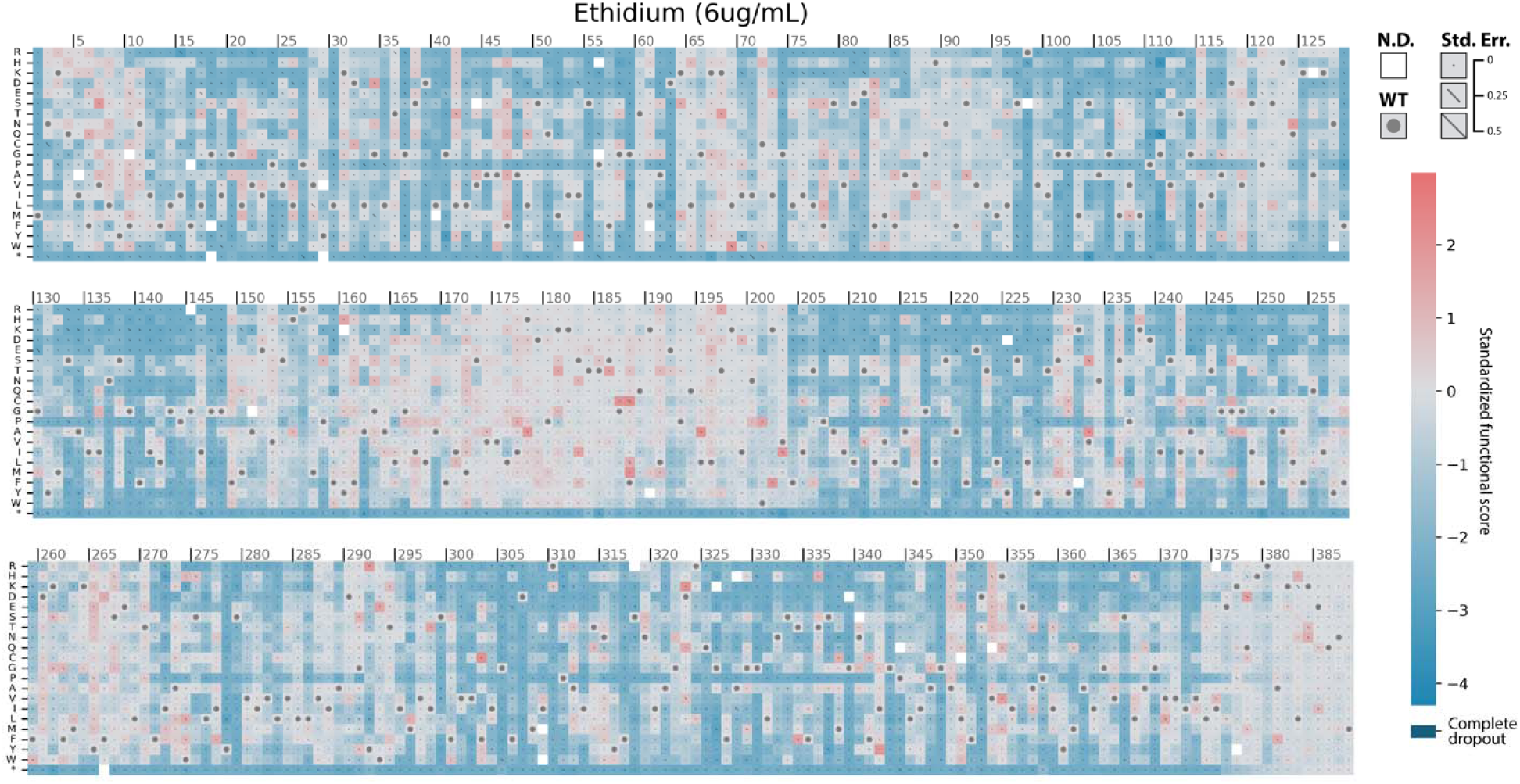
Sequence-function landscape of ethidium transport. Functional scores for all variants after 12 hours of competitive growth in the presence of 6μg/mL ethidium. The wild-type residue at each position is marked with a dot. Error values are shown as diagonal lines across each square, scaled so that the entire diagonal represents an error of 0.5 standardized functional score units. Variants with error > 0.5 are omitted (white). Variants which dropped out completely (zero observations after selection) are shown in dark blue.

**Supplementary Figure 5.**
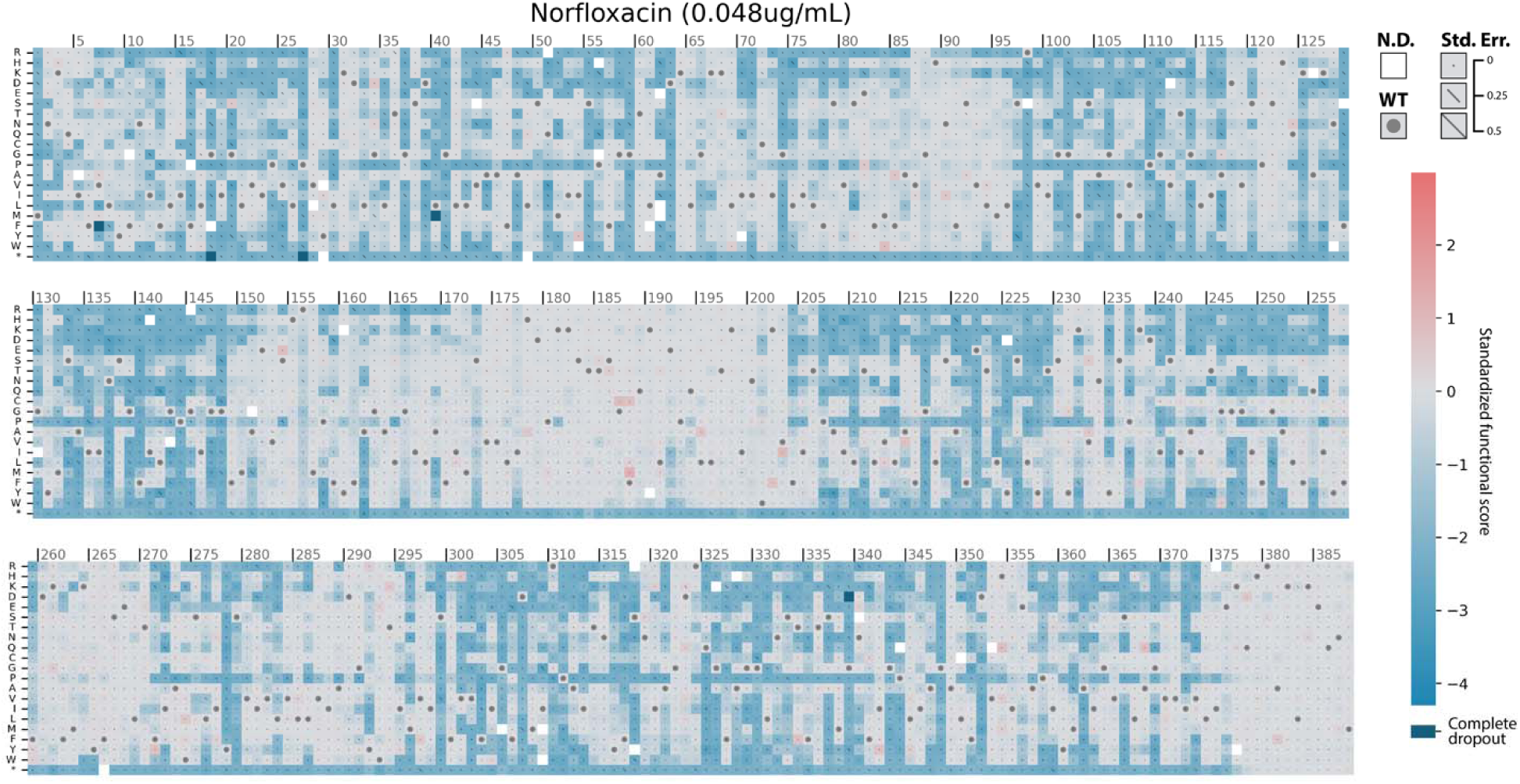
Sequence-function landscape of norfloxacin transport. Functional scores for all variants after 12 hours of competitive growth in the presence of 0.048μg/mL norfloxacin. The wild-type residue at each position is marked with a dot. Error values are shown as diagonal lines across each square, scaled so that the entire diagonal represents an error of 0.5 standardized functional score units. Variants with error > 0.5 are omitted (white). Variants which dropped out completely (zero observations after selection) are shown in dark blue.

**Supplementary Figure 6.**
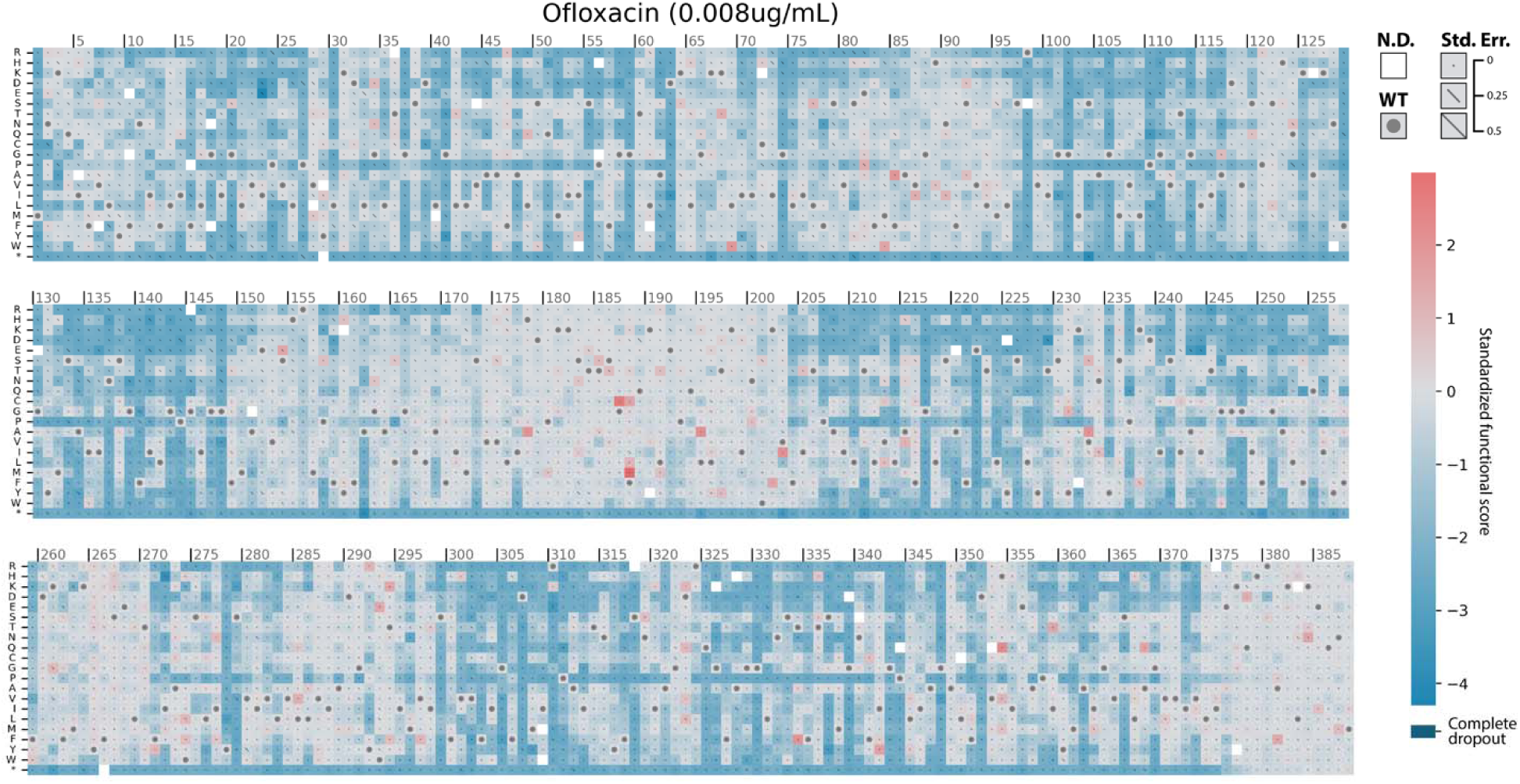
Sequence-function landscape of ofloxacin transport. Functional scores for all variants after 12 hours of competitive growth in the presence of 0.008μg/mL ofloxacin. The wild-type residue at each position is marked with a dot. Error values are shown as diagonal lines across each square, scaled so that the entire diagonal represents an error of 0.5 standardized functional score units. Variants with error > 0.5 are omitted (white). Variants which dropped out completely (zero observations after selection) are shown in dark blue.

**Supplementary Figure 7.**
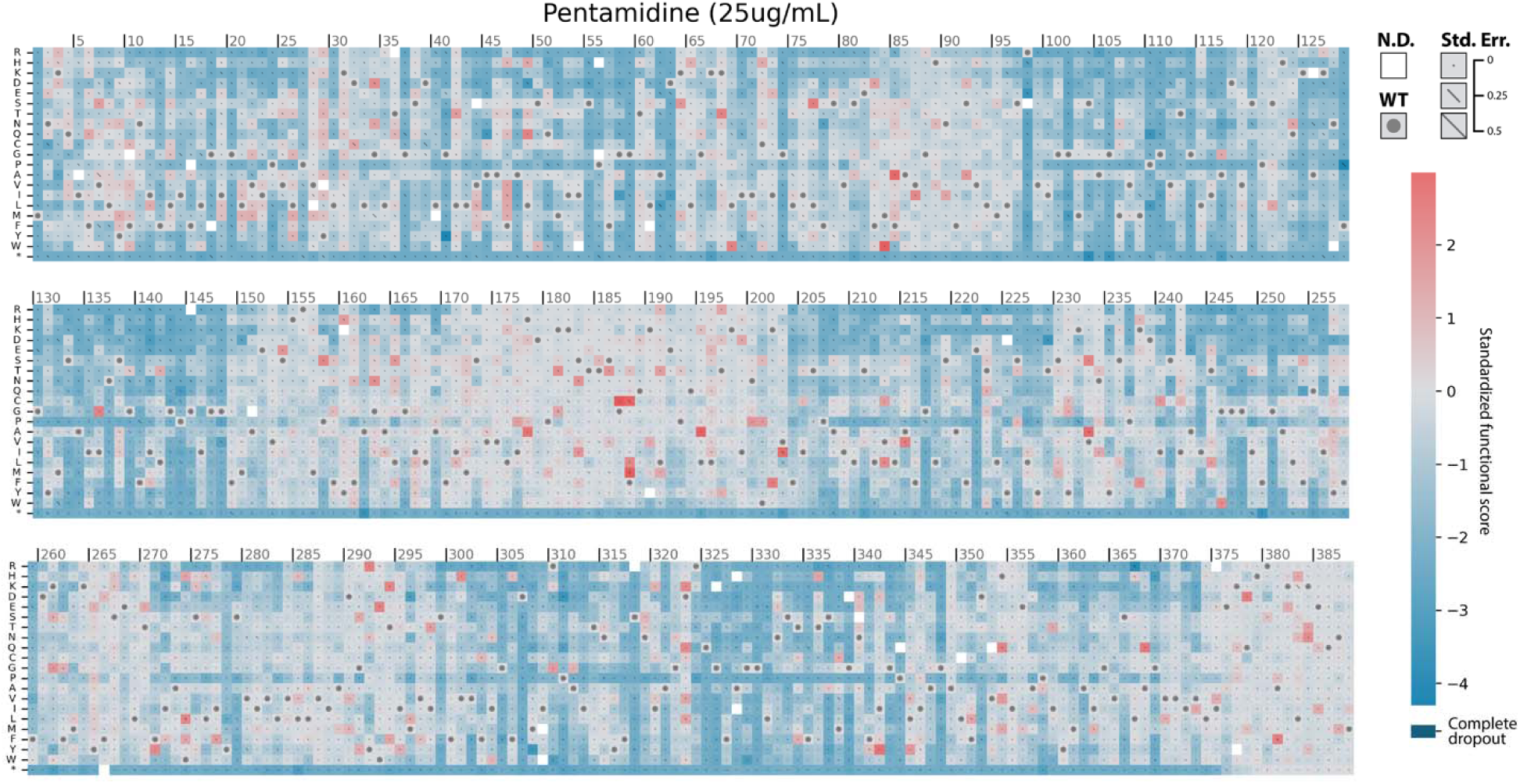
Sequence-function landscape of pentamidine transport. Functional scores for all variants after 12 hours of competitive growth in the presence of 25μg/mL pentamidine. The wild-type residue at each position is marked with a dot. Error values are shown as diagonal lines across each square, scaled so that the entire diagonal represents an error of 0.5 standardized functional score units. Variants with error > 0.5 are omitted (white). Variants which dropped out completely (zero observations after selection) are shown in dark blue.

**Supplementary Figure 8.**
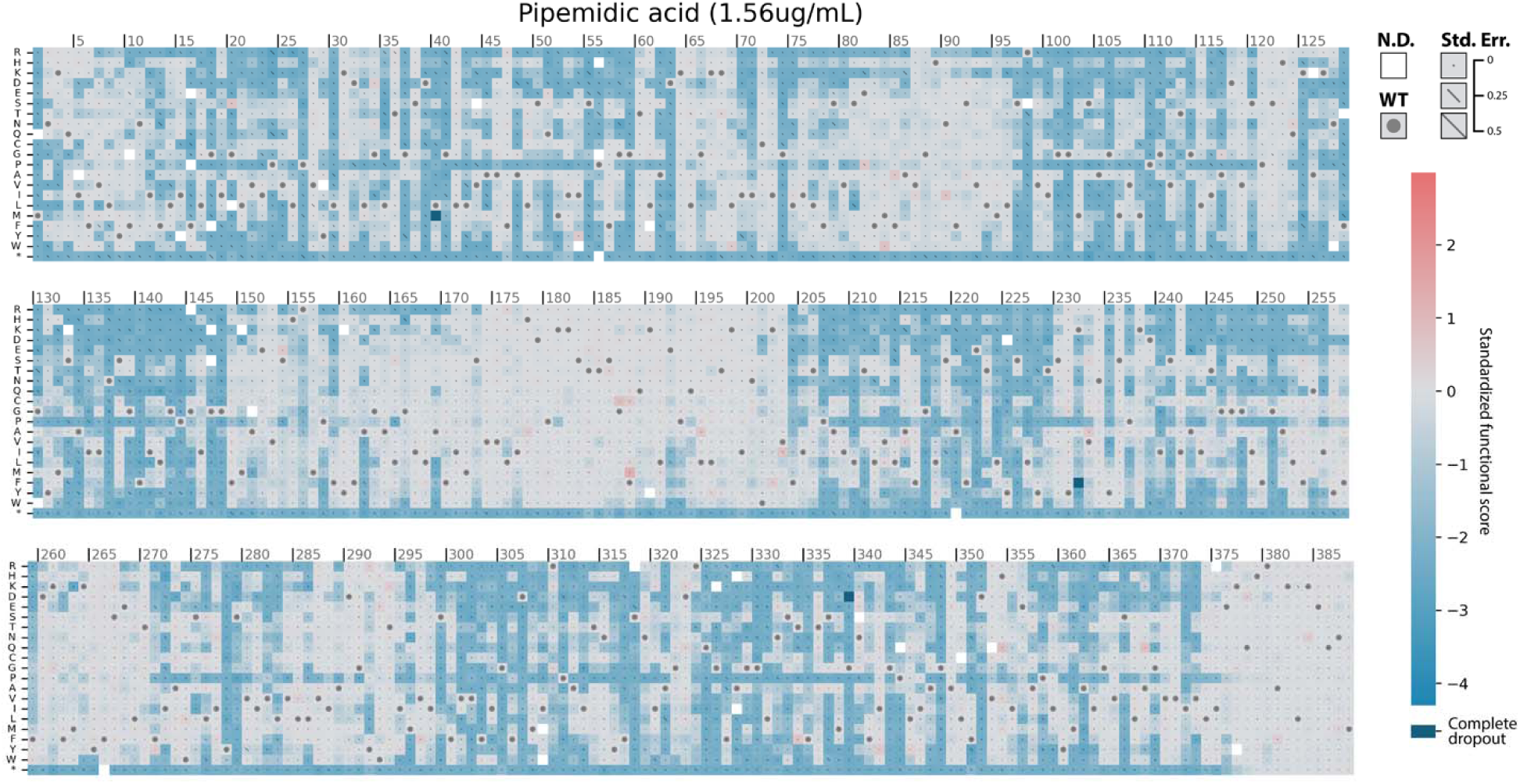
Sequence-function landscape of pipemidic acid transport. Functional scores for all variants after 12 hours of competitive growth in the presence of 1.56μg/mL pipemidic acid. The wild-type residue at each position is marked with a dot. Error values are shown as diagonal lines across each square, scaled so that the entire diagonal represents an error of 0.5 standardized functional score units. Variants with error > 0.5 are omitted (white). Variants which dropped out completely (zero observations after selection) are shown in dark blue.

**Supplementary Figure 9.**
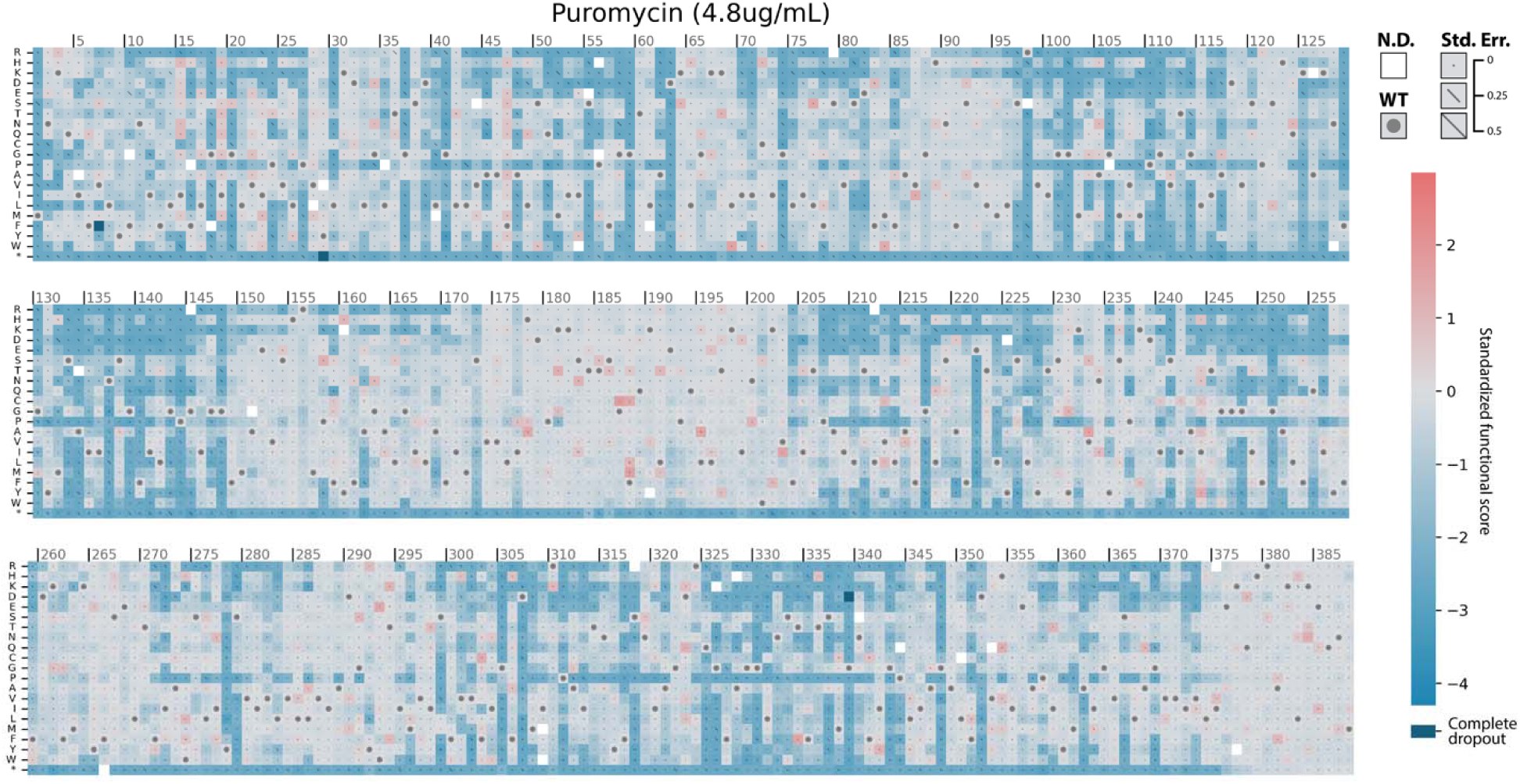
Sequence-function landscape of puromycin transport. Functional scores for all variants after 12 hours of competitive growth in the presence of 4.8μg/mL puromycin. The wild-type residue at each position is marked with a dot. Error values are shown as diagonal lines across each square, scaled so that the entire diagonal represents an error of 0.5 standardized functional score units. Variants with error > 0.5 are omitted (white). Variants which dropped out completely (zero observations after selection) are shown in dark blue.

**Supplementary Figure 10.**
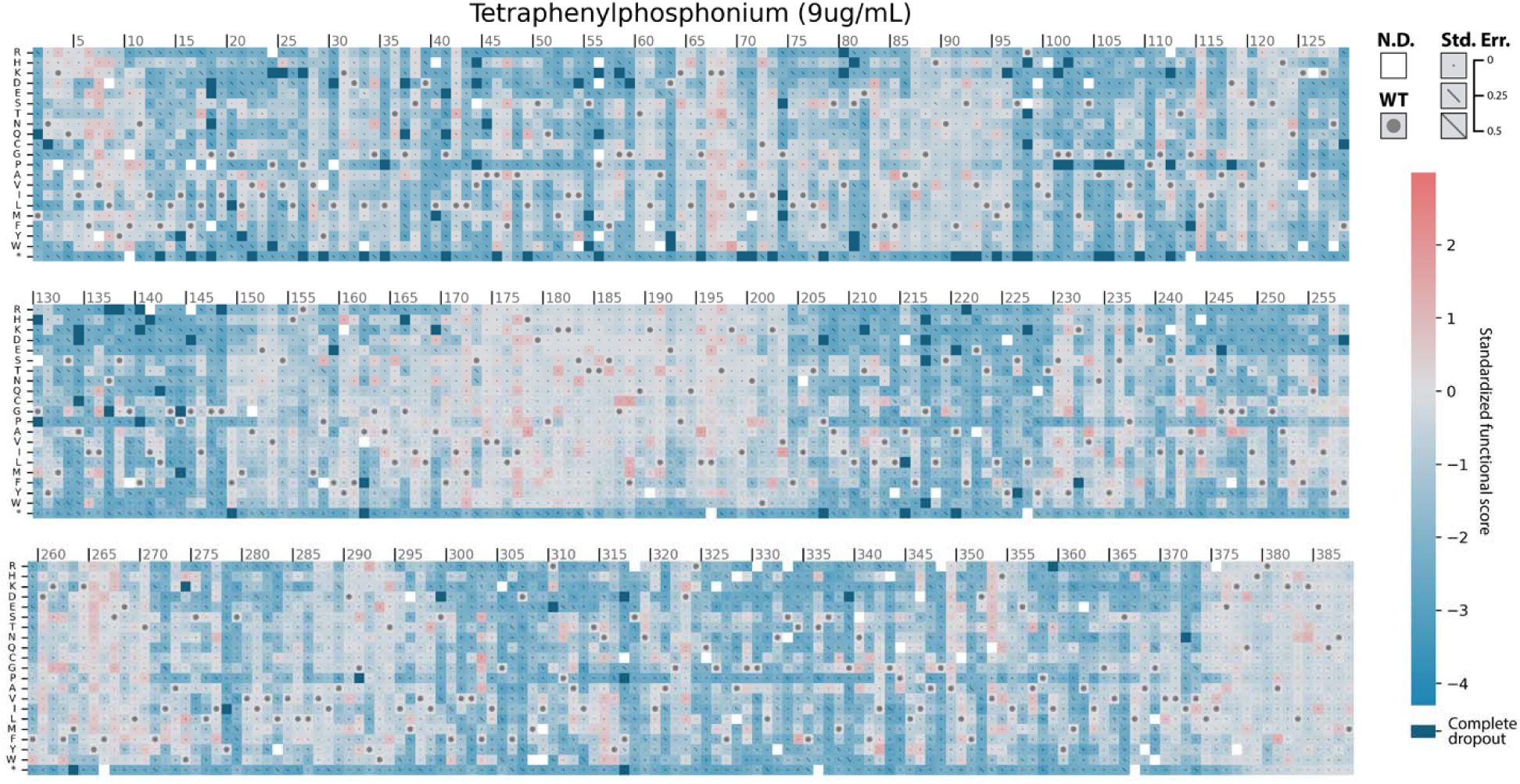
Sequence-function landscape of tetraphenylphosphonium transport. Functional scores for all variants after 12 hours of competitive growth in the presence of 9μg/mL tetraphenylphosphonium. The wild-type residue at each position is marked with a dot. Error values are shown as diagonal lines across each square, scaled so that the entire diagonal represents an error of 0.5 standardized functional score units. Variants with error > 0.5 are omitted (white). Variants which dropped out completely (zero observations after selection) are shown in dark blue.

**Supplementary Table 1.**
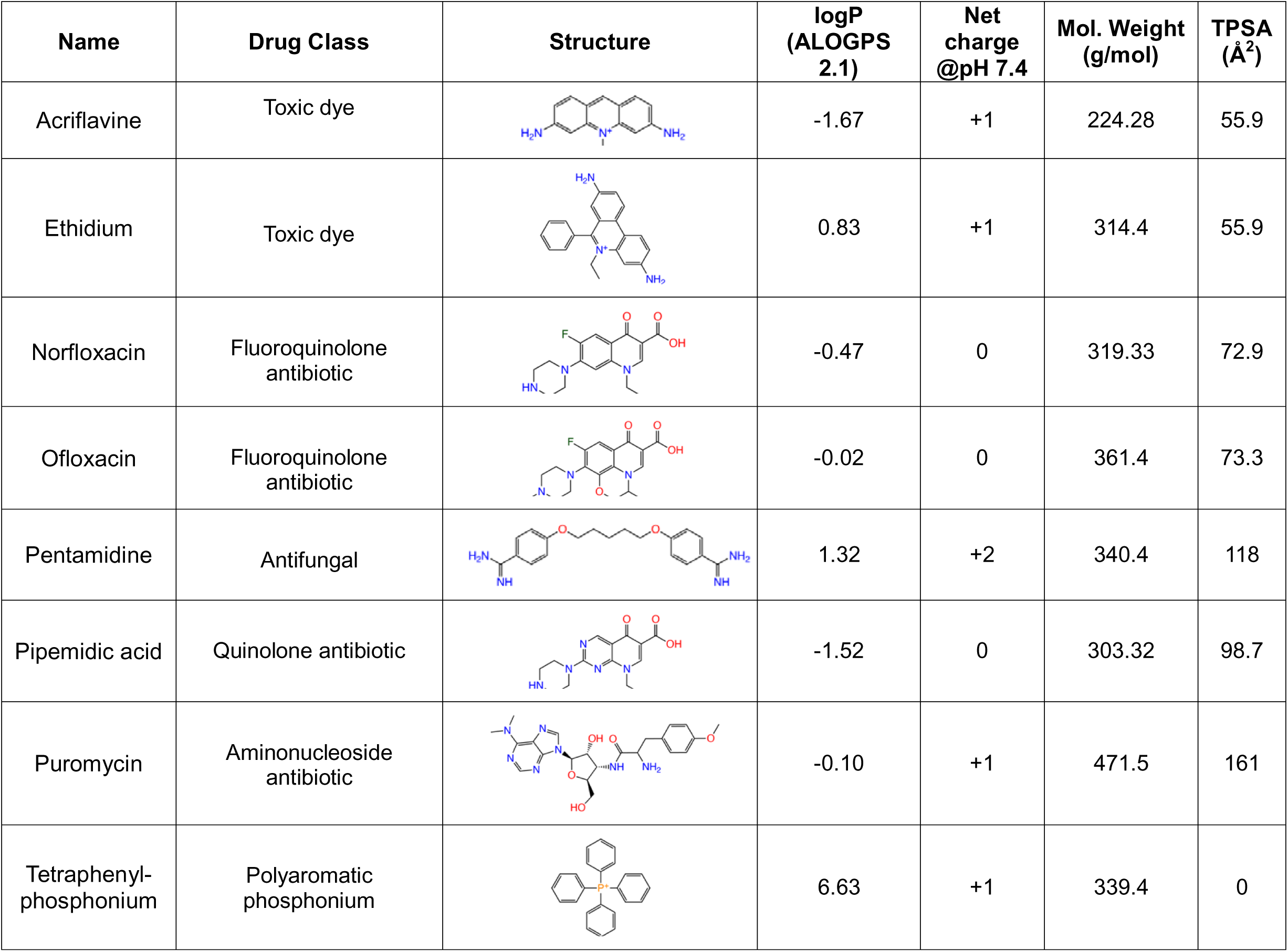
Substrate chemical properties. Chemical properties of the substrates used for selection. Values are shown for hydrophobicity (logP), net charge at physiological pH, molecular weight (g/mol), and topological polar surface area (TPSA, Å^2^). logP values were calculated using ALOGPS 2.1^89^. Physiological charge was retrieved from DrugBank^90^. TPSA was retrieved from DrugBank (acriflavine) or PubChem^91^ (all other substrates).

| Name | Drug Class | Structure | logP<br>(ALOGPS<br>2.1) | Net<br>charge<br>@pH 7.4 | Mol. Weight<br>(g/mol) | TPSA<br>(Å<sup>2</sup>) |
| --- | --- | --- | --- | --- | --- | --- |
| Acriflavine | Toxic dye |  | -1.67 | +1 | 224.28 | 55.9 |
| Ethidium | Toxic dye |  | 0.83 | +1 | 314.4 | 55.9 |
| Norfloxacin | Fluoroquinolone<br>antibiotic |  | -0.47 | 0 | 319.33 | 72.9 |
| Ofloxacin | Fluoroquinolone<br>antibiotic |  | -0.02 | 0 | 361.4 | 73.3 |
| Pentamidine | Antifungal |  | 1.32 | +2 | 340.4 | 118 |
| Pipemidic acid | Quinolone antibiotic |  | -1.52 | 0 | 303.32 | 98.7 |
| Puromycin | Aminonucleoside<br>antibiotic |  | -0.10 | +1 | 471.5 | 161 |
| Tetraphenyl-phosphonium | Polyaromatic<br>phosphonium |  | 6.63 | +1 | 339.4 | 0 |

**Supplementary Figure 11.**
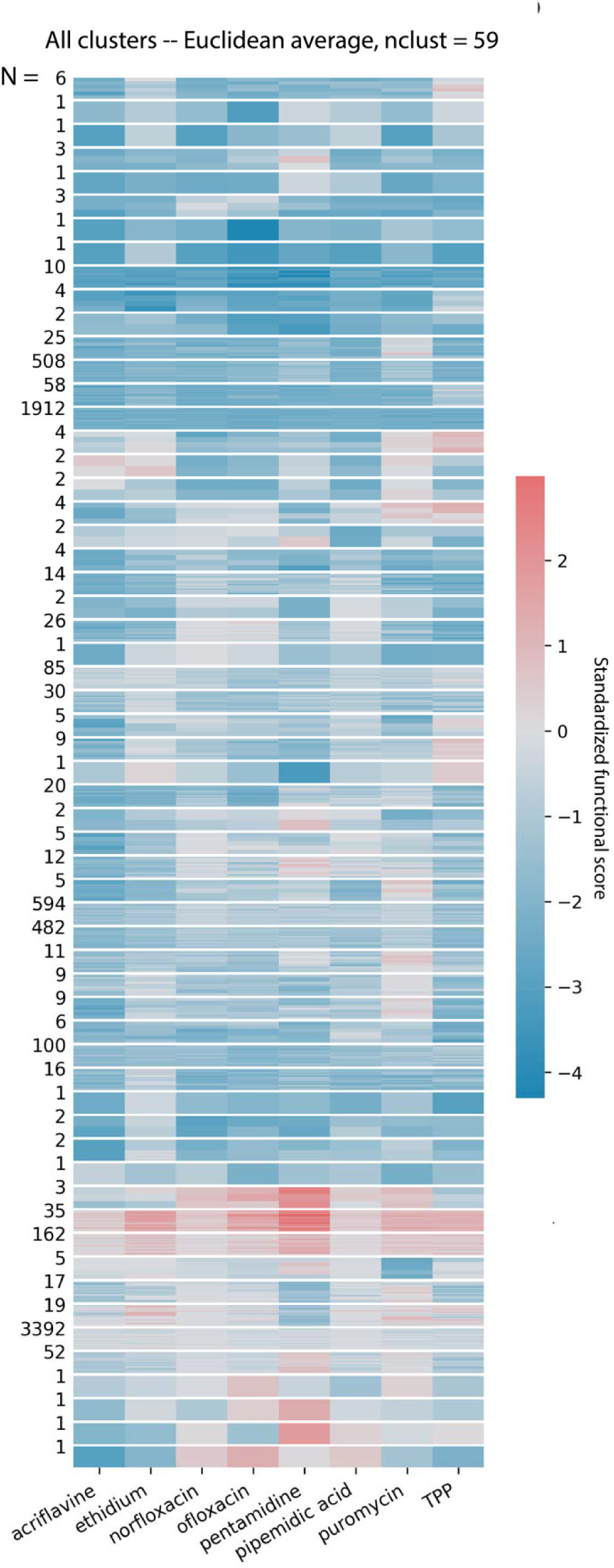
Unsupervised hierarchical clustering. All clusters observed after unsupervised hierarchical clustering with the Euclidean distance metric and average linkage method. The dendrogram was cut to yield 59 clusters based on gap statistic and domain knowledge.

**Supplementary Figure 12.**
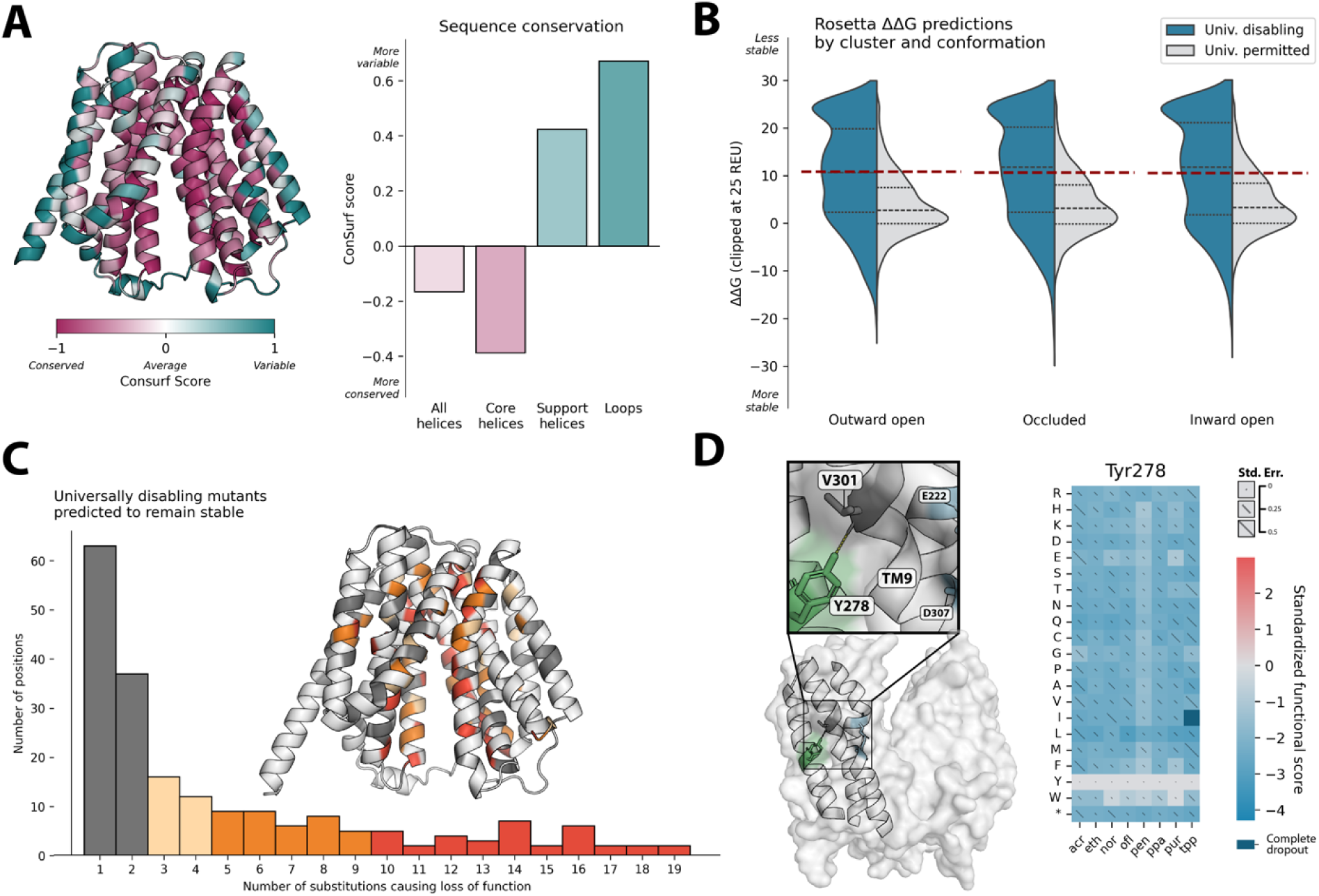
Universal functional hotspots. [A] NorA sequence conservation calculated by ConSurf and projected onto the NorA structure (AlphaFold). Average conservation scores for all helices, core helices only (TMs 1, 2, 4, 5, 7, 8, 10, 11), support helices only (TMs 3, 6, 9, 12), and loops shown in bar plot. [B] Split violin plot showing the distribution of Rosetta predicted ΔΔG_mut_ for variants found in the universally disabling cluster and universally permitted cluster in each conformation tested. Misfolding cutoff (1 standard deviation) is shown by red dotted lines. [C] Variants from the universally disabling cluster that are not predicted to misfold by Rosetta. Positions that cause complete loss of function without destabilizing the protein for at least 5 of the 19 possible substitutions are considered functional hotspots and are shown in orange and red. Complete list these positions can be found in Supplementary Table 2. [D] Single-residue heatmap showing universal depletion of Y278, a novel functional hotspot identified here, with potential mechanism highlighted on the NorA structure. Abbreviations: acr (acriflavine); eth (ethidium); nor (norfloxacin); ofl (ofloxacin); pen (pentamidine); ppa (pipemidic acid); pur (puromycin); tpp (tetraphenylphosphonium).

**Supplementary Table 2.** Universal functional hotspots. Residues for which at least 5 substitutions result in complete loss of function for all substrates while remaining stable in at least one conformation (orange and red positions in Supplementary Figure 12C). Positions in bold result in loss of function without destabilization for 10+ missense mutations (red in Supplementary Figure 12C).

| **Function** | **Residues** | **Citation** |
| --- | --- | --- |
| Proton binding/coupling mechanisms | **N137, E222, D307**<br>Y278 | 34; This study (Y378)4/10/2025 4:41:00 PM |
| Gating conformational change | **F129, S318, Q325,**<br>K125 | 34 |
| Motif D2 "lgxxxxxPvxP" | **G18, G20, P24, P27** | 43, 44, 46 |
| Motif A "GxLaDrxGrkxx(x)I" | **G59, D63,**<br>A62 | 43, 44, 46 |
| Motif B "lxxxRxxqGxgaa" | **R98,**<br>A105, G106 | 43, 44, 46 |
| Motif C "gxxxGPxxGGxl" | **P144, G148,**<br>F140, I141, G145, G147 | 43, 44, 46 |
| Motif G "GxxxGPL" | G339, P344 |  |
| Helix-helix interfaces | **G37, G41, A45, A48, S55, G114, A117, A134, G217, S241, G248, S279, G305, G330, S337, G348,**<br>G58, G74, G130, S133, A162, A252, S299, G326, A328, G329, A362, S366 | This study; See ref. 47 |
| Extracellular rim | **L30,**<br>D39, S226, L227, P237, D352 | This study; See ref. 51 |
| Conformational hinge points | M52, T245, Q255 | This study; See ref. 54 |
| Helix kinking | **P110, P311,**<br>P56 | This study; See ref. 52, 53 |
| General substrate / inhibitor recognition | Q51, T211, N332 | This study; See ref. 55–58 |
| Expression | **M1,**<br>N2 | See ref. 92 |

**Supplementary Figure 13:**
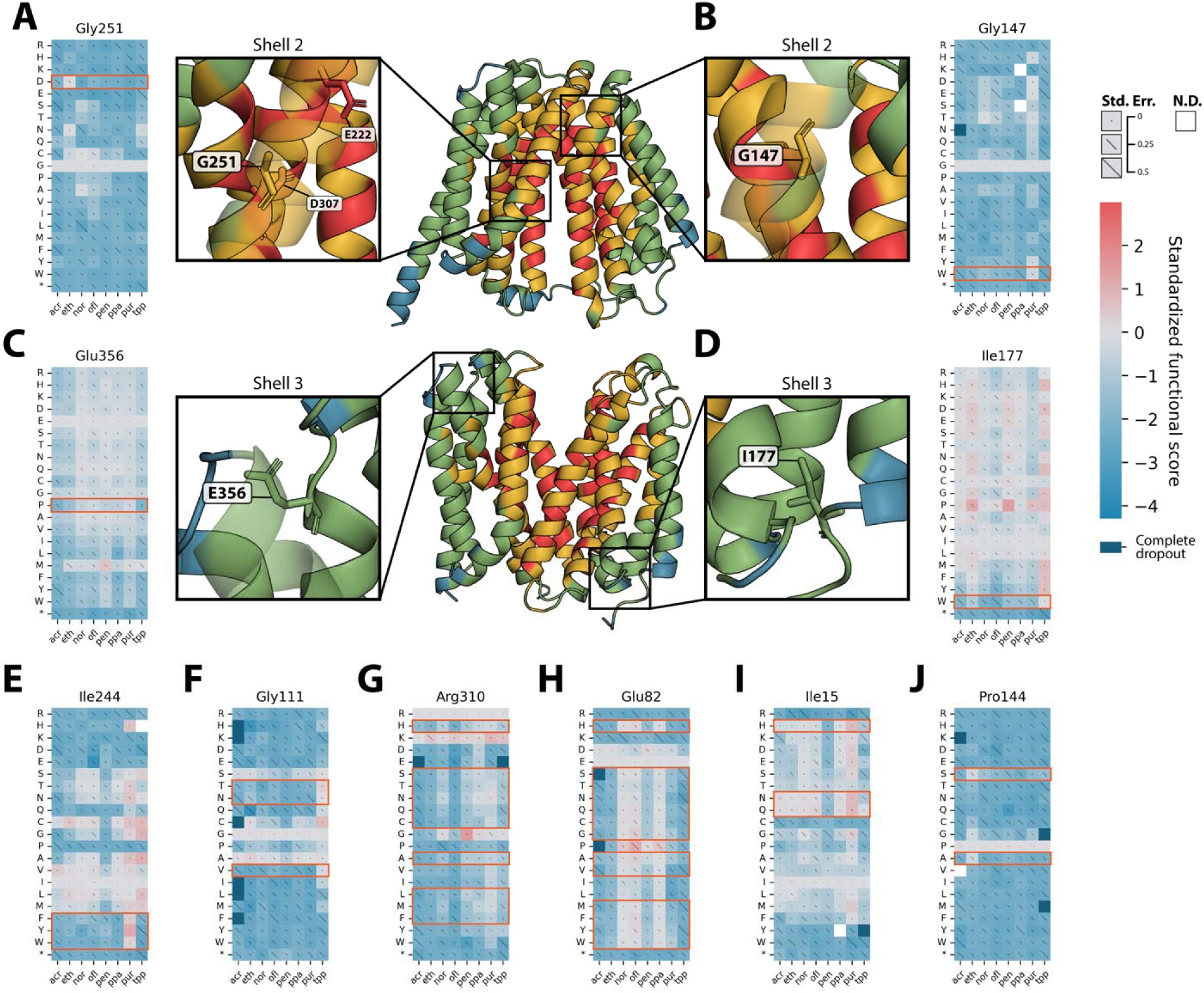
Single-position heatmaps for select specificity-driving residues. [A] Heatmap for Gly251, a Shell 2 residue which allows ethidium transport only when mutated to Asp. [B] Heatmap for Gly147, a Shell 2 residue which allows puromycin transport only when mutated to Trp. [C] Heatmap for Glu256, a Shell 3 residue which allows polyaromatic cation transport only when mutated to Pro. [D] Heatmap for Ile177, a Shell 3 residue which allows TPP transport only when mutated to Trp. [E-J] Single-residue heatmaps for select residues represented in Fig. 3B-G. Mutations belonging to the cluster discussed in the text are highlighted in with orange boxes. Abbreviations: acr (acriflavine); eth (ethidium); nor (norfloxacin); ofl (ofloxacin); pen (pentamidine); ppa (pipemidic acid); pur (puromycin); tpp (tetraphenylphosphonium).

**Supplementary Figure 14.**
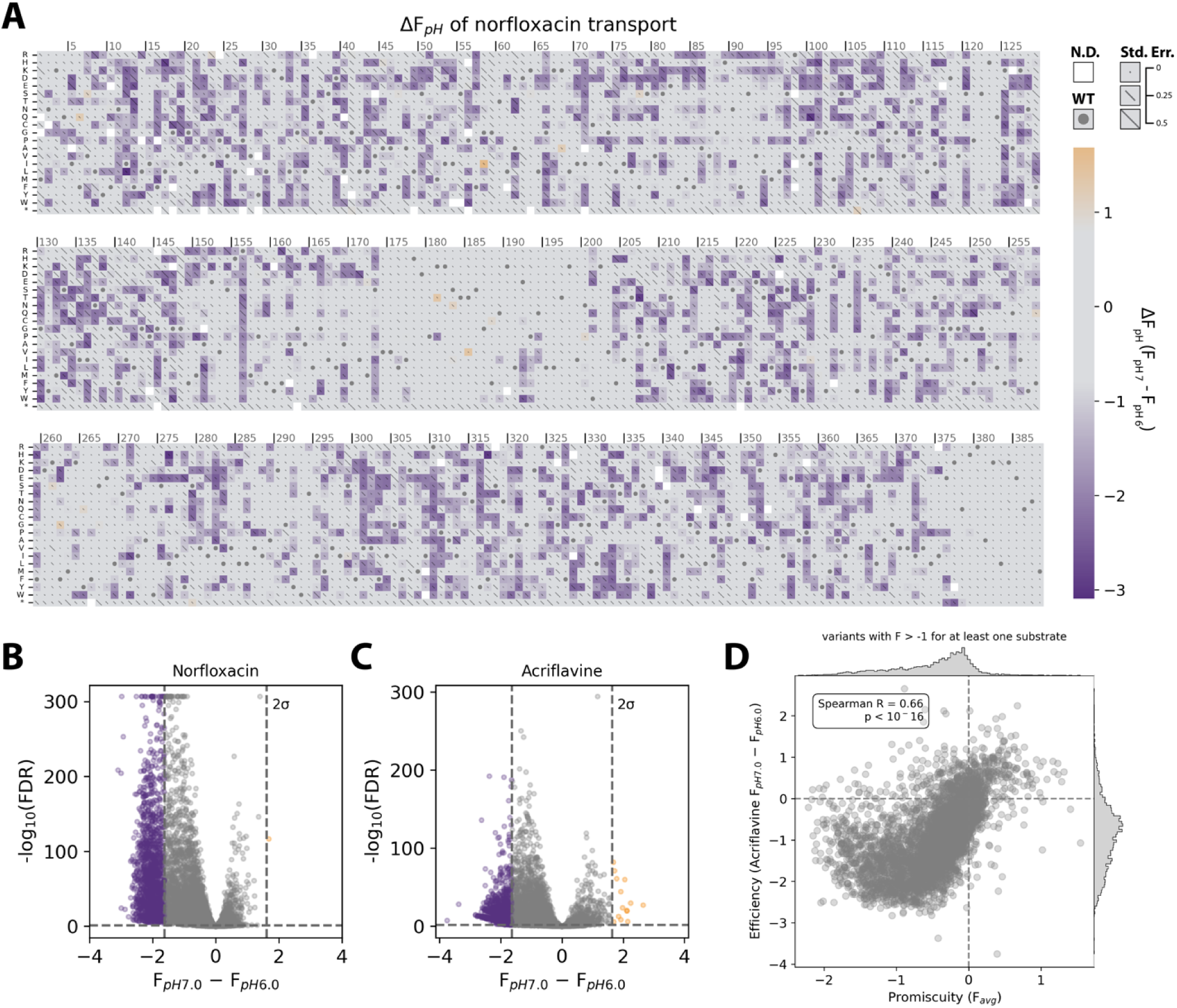
Sequence determinants of pH sensitivity. [A] Norfloxacin-based ΔF_pH_ scores for all variants measured in the NorA DMS library. The wild-type residue at each position is marked with a dot. Propagated error values are shown as diagonal lines across each square, scaled so that the entire diagonal represents an error 0.5 standardized functional score units. Variants with error > 0.5 and variants which dropped out of selection completely in either pH condition are not shown. [B] Volcano plot showing significance and magnitude of norfloxacin-based pH sensitivity changes for all variants. [C] Volcano plot showing significance and magnitude of acriflavine-based pH sensitivity changes for all variants. [D] Scatter plot comparing promiscuity (average functional score in all specificity screens) with efficiency (ΔF_pH_ from acriflavine pH-sensitivity screen) for variants with activity on at least one substrate.

**Supplementary Figure 15.**
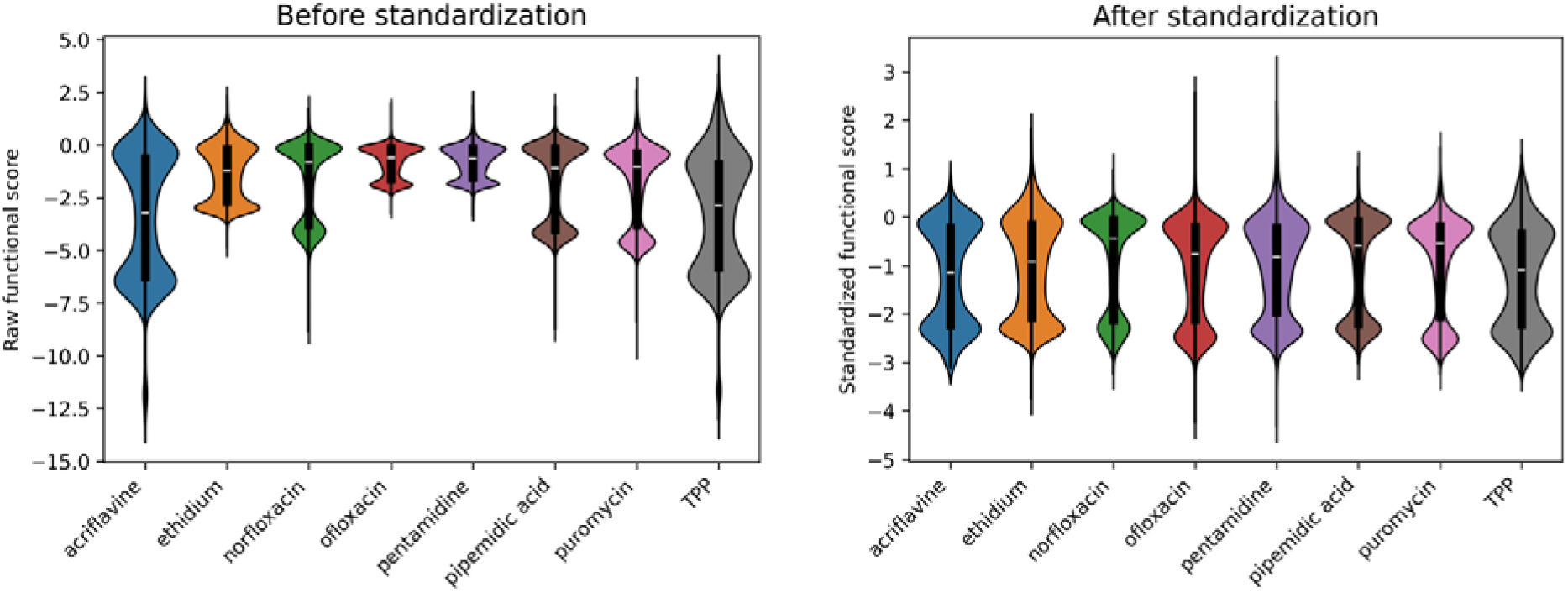
Standardization of functional scores. Violin plots demonstrating how functional score standardization allows direct comparison of scores from different substrate selections. Scores from each selection were divided by their own standard deviation (see Methods).

## Methods

### Creation of MG1655-ΔacrAB host strain

*In vivo* transport experiments can be confounded by the host’s native efflux pumps, which often have overlapping specificity profiles. To mitigate this, we used lambda red recombinase^93^ to create the drug-hypersensitive *E. coli* strain MG1655-ΔacrAB. This strain is deleted of *E. coli*’s primary efflux machinery and exhibits significantly increased susceptibility to most antimicrobials tested without significant growth defects in the absence of drug^39,40^. By expressing NorA variants in this genetic background, small changes in drug resistance are more easily observed^42^.

A DNA fragment containing a kanamycin resistance cassette flanked by Flp recombinase recognition target (FRT) sites and at least 50bp of homology for sequence up– and down-stream of the acrA and acrB genes was created by PCR amplification with KAPA HiFi polymerase (Roche) and primers synthesized by IDT. Homology arms were incorporated as 5’ primer “tails.” The resistance cassette and FRT sites were amplified using colony PCR on strain of *E. coli* from the KEIO collection^94^.

A 25mL culture of MG1655 *E. coli* harboring the arabinose-inducible temperature-sensitive lambda red recombinase plasmid pKD46 (Genbank AY048746) was induced with 50mM L-arabinose at OD600 0.1, growing at 30C. At OD600 0.6, electrocompetent cells were prepared by pelleting and washing cells with 10% glycerol 3 times, finally resuspending in 50μL 10% glycerol. 25μL of electrocompetent MG1655+pKD46 was transformed by electroporation with 500ng of the kanamycin resistance cassette DNA fragment prepared previously. After 2 hours of recovery at 37C, cells were plated on LB agar containing kanamycin and grown at 37C.

A kanamycin-resistant clone was confirmed to be missing the acrAB genes by colony PCR. This clone was transformed with an arabinose-inducible temperature-sensitive Flp recombinase plasmid (AddGene Plasmid #122969)^95^ and induced with 100mM L-arabinose for 8 hours at 30C. Culture was diluted and plated on LB agar containing no antibiotic and grown at 37C to cure the Flp recombinase plasmid. Removal of the kanamycin cassette was confirmed by colony PCR and streaking on agar plates with 30μg/mL kanamycin. Curing of the lambda red and Flp recombinase plasmids was confirmed by streaking on agar plates with 100μg/mL carbenicillin. The resulting MG1655-ΔacrAB *E. coli* was used as the host background for all experiments described herein.

### Library construction

We built a wild-type NorA expression plasmid which was used as a template for library construction. We obtained the most active norA allele^45^, norAII, by colony PCR of wild-type *S. aureus* and used Gibson assembly to clone this gene into a medium copy plasmid backbone (pBR322 origin with *rop*) carrying resistance to kanamycin, which is not a substrate of NorA. The norA gene is driven by the weak constitutive Anderson promoter pJ23114. This plasmid and all plasmids described here were sequence-verified with whole-plasmid sequencing (PlasmidSaurus).

Oligonucleotides encoding each NorA variant were synthesized by Twist Bioscience. The NorA variant library was designed by replacing all wild-type residues to all other amino acids or a stop codon, using the most common *E. coli* codon possible without introducing interfering restriction sites. Due to DNA synthesis length limitations, oligonucleotides were organized into five subpools spanning the ∼1.2kb gene: Residues 1-72, 73-154, 155-231, 232-311, and 312-388. Within each synthesized oligonucleotide, the NorA variant sequence was flanked by BsaI restriction sites, as well as forward and reverse priming sequences for amplifying each subpool from the oligonucleotide pool independently.

For each subpool, a “backbone” fragment was amplified from the wild-type NorA plasmid using primers that added BsaI restriction sites and did not include any norA sequence downstream of where the subpool begins. The reverse primer also added a DNA “barcode” containing 20 random nucleotides to facilitate identification of variants by short-read sequencing. With the exception of the 3’-most final subpool, a separate fragment was amplified containing the wild-type norA sequence downstream of where that subpool ends. Subpools 1-4 were assembled with three-part Golden Gate assemblies using 50fmol of backbone and 100fmol of insert fragments (BsaI-HF NEBridge Golden Gate Assembly kit, NEB). The 3’-most subpool 5 was assembled using a two-part Golden Gate reaction because the oligonucleotide subpool extended to the very end of the norA gene. (Supplementary Fig. 2D).

All fragments were digested with DpnI (NEB) prior to Golden Gate assembly. To prevent persistent backbone re-ligation, the backbone fragment for subpool 2 was also digested with BsaI-HF (NEB) and Antarctic phosphatase (NEB) prior to Golden Gate assembly. Golden Gate reactions were performed using 30 cycles of [37C for 5 minutes, 16C for 5 minutes], followed by a 5-minute 60C inactivation. After assembly, each Golden Gate reaction was dialyzed for 1 hour with water on silica membranes (MF-Millipore Membrane Filter, 0.025µm pores, Millipore Sigma) before transforming 5μL dialyzed Golden Gate assembly into 50μL electrocompetent DH10B *E. coli* prepared in-house as previously described. After a 1-hour recovery in SOC medium at 37C, cells were spread on LB agar plates with 30μg/mL kanamycin to assess transformation efficiency and diluted in fresh liquid cultures to intentionally bottleneck the library, such that the number of barcodes per variant would be 25-30. After 12 hours of growth, libraries were miniprepped and stored at –20C.

DNA from each subpool miniprep was proportionally mixed according to the number of variants in each subpool. 200ng of this mixture was transformed into 100μL of electrocompetent MG1655-ΔacrAB. This complete norA library was allowed to recover at 37C in SOC medium for 1 hour before diluting 1:50 in 100mL LB containing 30μg/mL kanamycin and growing at 37C until the OD600 read 1.5 (about 4.5 hours from dilution). Culture was mixed 1:1 with 50% glycerol and stocked in 2mL aliquots at –80C.

### Barcode mapping

To map barcodes to individual variants using short-read Illumina sequencing, “mapping constructs” were created to position barcodes directly adjacent to the variable region of the NorA gene. Using PCR and Golden Gate assembly, we removed DNA from subpools 1-4 and reassembled the plasmid such that each subpools’ variable region and barcode region could be sequenced in the same 2×300 paired-end Illumina sequencing read. These PCR reactions used 10ng of template and 12 cycles, the minimum number of PCR cycles necessary to obtain enough product (in order to minimize introduction of template switching and PCR bias). No mapping construct was necessary for the 3’-most subpool 5, because the variable region is already adjacent to the barcodes. (Supplementary Fig. 1A)

To determine which barcode mappings were of high enough quality to be reliable for future experiments, we employed a read-count cutoff and chastity filter. The read count cutoff was set individually for each subpool at the elbow of the curve comparing read cutoff to number of remaining sequences (minimum 5, maximum 10). For each of the remaining barcodes, a chastity value was calculated by dividing the number of counts for the most common barcode-variant mapping by the sum of the top two most common barcode-variant mappings. Barcodes with a chastity value of less than 0.975 were removed. Quality filtering for a representative subpool is shown in Supplementary Fig. 1B and C.

To empirically confirm that our barcode mapping and quality filtering methods were sufficiently stringent, we performed growth selections on an individual subpool. Because the variable region of each subpool spans only about 240bp, Illumina sequencing can capture variants either by barcode or by direct variant sequencing. We found that functional scores calculated by counting barcodes correlated very well with those by direct variant sequencing (Supplementary Fig. 1D and E; norfloxacin selection: R^2^ = 0.99; acriflavine selection: R^2^ = 0.98), indicating that sequencing barcodes recapitulates the same information as the gold-standard of direct variant sequencing.

### Selection of drug toxicity growth conditions

The eight NorA substrates used in this study were identified from the literature and confirmed to be reproducible substrates in our hands (acriflavine, ethidium bromide, norfloxacin, ofloxacin, puromycin, and pipemidic acid were obtained from Sigma-Aldrich. Pentamidine isethionate salt and tetraphenylphosphonium bromide are from Fisher Scientific). We used broth microdilution to determine the minimum inhibitory concentration (MIC) of each drug^96^ for MG1655-ΔacrAB *E. coli* expressing wild-type NorA or the catalytically inactive variant of NorA E222A. For each drug, we chose a selection concentration between the MIC and the IC_50_ to allow for dynamic range to observe variants that perform better or worse than wild type. A relatively short selection time of 12 hours was chosen to minimize the risk for suppressor mutations arising in the library.

### Performing drug toxicity library selections

Each drug toxicity selection was performed according to the following protocol. A 2mL aliquot of glycerol-stocked NorA library was retrieved from storage at –80C and allowed to thaw completely on ice. Each milliliter of this stock contains enough colony-forming units (CFU) to ensure 500–1000X coverage of every barcode in the library, accounting for 60–70% cell death from freeze-thaw. All following steps are performed in duplicate: 1mL of this stock was diluted 1:10 into LB medium containing 30μg/mL kanamycin and allowed to recover at 37C with shaking until an OD600 measurement of 0.8-1.0 (about 2 hours). To remove dead cells and glycerol, the starter culture was diluted 1:200 in fresh LB medium with 30μg/mL kanamycin (150μL starter culture in 30mL fresh medium). This “input culture” was allowed to grow at 37C with shaking to an OD600 of 0.5 (about 3 hours). A plasmid miniprep was performed using 10mL of input culture to isolate DNA for “pre-selection” frequencies of library variants.

100mL of LB medium with 30μg/mL kanamycin was prepared with the appropriate concentration of selection drug in a 250mL flask. The selection culture was seeded with 125μL of OD 0.5 input culture to achieve roughly the standard antimicrobial testing inoculum size of 5×10^5^ CFU/mL while maintaining 200-400X coverage of all barcodes. Selection cultures were grown at 37C with shaking for 12 hours, then miniprepped to isolate DNA for “post-selection” frequencies of library variants.

### Library preparation and sequencing

For both replicates of input and selected libraries, the region of the NorA library containing barcodes was amplified by PCR using primers that attach Illumina sequencing primer binding sites. This PCR used 1ng of template with a total of 12 PCR cycles. After purifying PCR product, a second PCR was performed to attach i5 and i7 indexes and P5 and P7 sequences. The second PCR used 10ng template with a total of 8 PCR cycles. The minimum number of cycles necessary to obtain enough DNA was used in both PCR steps to minimize introduction of PCR bias and template switching.

After confirming fragment size and purity by gel electrophoresis and measuring DNA concentration with a Qubit fluorimeter (Thermo Fisher Scientific), libraries were pooled and prepared for sequencing on the Illumina NextSeq 1000 using a 750pM library concentration and 5-15% PhiX spike-in to account for low base diversity. When sequencing, we targeted at least 250X coverage per-variant. This coverage target is based on *in silico* coverage down-sampling which showed 250X as the point of diminishing returns for improving correlation between replicates and between barcode– and variant-derived functional scores.

### Sequencing data processing and calculation of functional scores

Raw fastq files were retrieved from the NextSeq 1000 and processed using compute resources at the UW Madison Biochemistry Department. First, paired-end reads were merged using PEAR with a minimum overlap of 10bp^97^. Merged reads were filtered for quality, requiring all bases in each read have a Phred score of at least Q20 and at least 95% of bases have a Phred score of at least Q30.

Merged reads passing all quality filters were analyzed using a custom python script that does the following. First, all barcodes are counted and summed to yield the total number of observations for each variant. Functional scores are calculated using a wild type–normalized log ratio of pre-to post-selection counts, based on that used in Enrich2 software^98^: F = ln((counts_var, selected_ + pseudocount)/counts_WT, selected_) – ln((counts_var, unselected_ + pseudocount)/counts_WT, unselected_). A pseudocount of 0.01 was used to allow calculation of scores for variants containing zero post-selection observations. Standard error for each score was calculated based on the method used in Enrich2: SE = sqrt((1/(counts_var, unselected_ + pseudocount)) + (1/(counts_WT, unselected_ + pseudocount)) + (1/(counts_var, selected_ + pseudocount)) + (1/(counts_WT, selected_ + pseudocount))). Functional scores and standard errors were calculated for each replicate selection separately, then merged using a robust maximum likelihood estimator as in Enrich2 using 100 iterations. To facilitate direct comparison of functional scores across substrates with varying potencies, a standardized functional score was also calculated by dividing each score and error by the standard deviation of all scores obtained for that substrate (Supplementary Fig. 15).

### Unsupervised hierarchical clustering

Agglomerative hierarchical clustering was performed using standardized functional scores with an arbitrary low score (–3) filled in for variants which dropped out of selection completely. Clustering distance metric (Euclidean) and linkage method (average) were chosen by measuring the cophenetic correlation and based on data type and characteristics. The number of clusters (59) was chosen to maximize the gap statistic and using domain knowledge.

All 59 clusters are shown in Supplementary Fig. 11. The 28 specificity-driving clusters shown in Figure 2 were identified procedurally as follows. First, clusters with only one or two members are removed. The remaining clusters are ranked by the Euclidean distance between the final two groups of *substrates* to be merged, normalized by the number of variants in that cluster. This is a measure of how well the mutations in each cluster separate different substrates into two groups—i.e., the magnitude of specificity shift for the given cluster. Clusters below the 25^th^ percentile by this metric are not shown in Figure 2.

### Calculation of specificity importance scores

Specificity importance (SI) scores were calculated using standardized functional scores with dropouts removed. Scores were first truncated with an upper maximum of zero to avoid high SI scores for variants with enhanced transport of some substrates but retaining nominal activity on other substrates. From this modified dataset, each variant’s minimum score was subtracted from the maximum score among all available values. To improve interpretability, a modified z-score was calculated form these range values by subtracting the median (0.68) and dividing by the standard deviation (0.52).

### Calculation of residue distances

Distances used in Figure 2F and Figure 4B were computed using the inward-open AlphaFold structure of NorA with the Biopython PDB package. “Shortest distance to [binding site residue / coupling residue]” refers to the minimum distance between any atom of the two residues being compared. Binding-site exposed residues were determined by examining the structure in PyMol: residues 12, 15, 16, 19, 20, 22, 23, 26, 40, 43, 44, 47, 48, 51, 55, 94, 98, 101, 105, 106, 109, 113, 126, 129, 132, 133, 136, 137, 140, 207, 211, 215, 218, 219, 222, 223, 241, 244, 248, 303, 306, 307, 310, 314, 329, 332, 333, 336, 337, 340, and 344. Residues with critical roles in coupling were determined from the literature: 98, 222, 307.

### Selection of pH stress growth conditions

A stock solution of bis-tris propane (Dot Scientific) buffer was prepared and adjusted to pH 6.0 or 7.0 and sterile filtered before adding to LB media to a final concentration of 75mM. These pH values were chosen because NorA-mediated resistance to norfloxacin was significantly reduced by pH 7.0 and bolstered by pH 6.0, while keeping the difference in the two pH values relatively minor to reduce the magnitude of interference by other confounding variables affected by pH.

### Calculation of pH sensitivity scores

Functional scores were calculated and standardized as described above for both pH 6.0 and pH 7.0 selections. The pH sensitivity score ΔF_pH_ was calculated as F_var, pH 7.0_ − F_var, pH 6.0_ and the error was propagated using the root sum of squared error of individual measurements.

### Clonal pH sensitivity measurements

Fourteen variants were built clonally with site-directed mutagenesis of the wild-type NorA plasmid (KLD Enzyme Mix, NEB) for validation of pH sensitivity of transport. Each variant and wild type was grown in the presence of varying concentrations of norfloxacin in 96-well plates at pH 6.0 and pH 7.0 in triplicate. After 16 hours of growth shaking at 37C, the OD600 was measured. IC_50_ values were calculated by fitting the curve of OD600 versus norfloxacin concentration to a four-parameter logistic function OD_min_ + (OD_max_-OD_min_) / (1 + ([drug] / IC_50_)^Hill^). The three replicate IC_50_ values and their errors were collapsed using a robust maximum likelihood estimator with 100 iterations.

A wild type-normalized “clonal functional score” was calculated for each variant at each pH by taking the log ratio of the variant’s IC_50_ to the wild-type IC_50_. A clonal pH sensitivity score was calculated by taking the difference between clonal functional scores at pH 7.0 and pH 6.0 and error was propagated.

### RT-qPCR

Triplicate overnight cultures of MG1655-ΔacrAB *E. coli* carrying plasmid expressing each of the clonally built validation variants were diluted 1:250 in 5mL fresh LB medium containing 30μg/mL kanamycin. Cultures were grown with shaking at 37C until an OD600 of 0.6. 2mL of culture was pelleted and a chloroform RNA extraction was performed. After 1 hour digestion with RNAse-free DNase I (Roche), RNA was cleaned and concentrated (RNA Clean & Concentrator-5, Zymo Research) before quantifying using Qubit fluorimeter.

qPCR reactions were prepared in triplicate using 400ng RNA template using the Luna Universal One-Step RT-qPCR kit (NEB). For each sample, two qPCR reactions were performed per replicate: one using primers to amplify a 105bp region of the NorA transcript which does not contain any of the validation variants, and one using primers to amplify a 105bp region of the cysG reference gene^99^. qPCR was performed at the UW Madison Biophysics Instrumentation Facility using a CFX Connect Real-Time PCR Detection System (Bio-Rad Laboratories).

A ΔCq was calculated by subtracting the cysG Cq from the norA Cq. The relative expression score for each variant is calculated as 2^-(ΔCq_var−ΔCq_WT)^. For each variant, the average and standard deviation of replicate relative expression scores were calculated.

### Rosetta ΔΔG predictions

Rosetta ΔΔG calculations were performed according to the protocol published in ref. 50 with slight modifications. Briefly, NorA structure files were downloaded from the RCSB protein databank and transformed into the membrane using the Orientations of Proteins in Membranes PPM 2.0 web server^100^. PDBs were cleaned of extraneous information, converted to Rosetta numbering, and chain of interest extracted using the clean_pdb.py script provided with the Rosetta software suite. A span file dictating membrane-spanning regions was generated using the Rosetta application spanfile_from_pdb^101,102^ and inspected for accuracy before continuing. The cleaned and transformed structure then underwent energy minimization using a cartesian fastrelax Rosetta scripts^103^ protocol adapted from ref. 50, using the following command: ${ROSETTASCRIPTS} –database ${ROSETTADB} –parser:protocol mp_cart_relax.xml – parser:script_vars repeats=5 energy_func=f19_cart_1.5.wts energy_fawtb=0 – in:file:s ${cleaned_pdb} –optimization::default_max_cycles 200 – mp:setup:spanfiles ${spanfile} –mp:scoring:hbond –mp:lipids:composition DLPC –mp::thickness 15 –relax:jump_move true –relax:coord_constrain_sidechains – relax:constrain_relax_to_start_coords –nstruct 1 –fa_max_dis 9.0 – ignore_unrecognized_res true –packing:pack_missing_sidechains false –ex1 –ex2 –flip_HNQ –missing_density_to_jump –score:weights f19_cart_1.5.wts –out:pdb – out:file:scorefile relax_scores.sc

The cleaned and relaxed structure file was then run through a complete ΔΔG scanning pipeline using the Rosetta cartesian_ddg^104^ application using the following command, with flags based on the cartesian_ddg docs and ref. 50: ${CARTESIAN_DDG} –database ${ROSETTADB} –s ${relaxed_pdb} –score:weights f19_cart_1.5.wts –in:membrane – mp:setup:spanfiles ${spanfile} –mp:lipids:composition DLPC –has_pore false – ddg:mut_file ${mutfile} –ddg:legacy true –ddg::dump_pdbs false –ddg:frag_nbrs 4 –ddg:optimize_proline true –ddg:cartesian –ddg:bbnbrs 1 –ddg:iterations 3 – force_iterations false –fa_max_dis 9.0 –missing_density_to_jump ΔΔG was calculated as ΔG_mut_ – ΔG_WT_ using the average energy from three simulations. This process was repeated for three experimentally solved conformations of NorA: outward-open (7LO8^4^), occluded, and inward-open (9B3L and 9B3M^35^).

## Acknowledgements

This material is based upon work supported in part by the Great Lakes Bioenergy Research Center, U.S. Department of Energy, Office of Science, Biological and Environmental Research Program under Award Number DE-SC0018409. This work was also supported by R35GM141748 to K.A.H.W. We would like to thank Dr. Max Frenkel for advice on DNA barcoding and quality filtering, Dr. Milo Lin for helpful conversations on transporter thermodynamics, and Dr. Ivan Rayment, Dr. Betty Craig, and Dr. Milo Lin for comments on the manuscript. We also thank current and former members of the Raman Lab, including Dr. Christian MacDonald, Dr. Tony Meger, James Corban, Dr. Phil Huss, Nate Novy, and Jonah Schwartz for discussions on data analysis and troubleshooting. Finally, we thank current and former members of the Henzler-Wildman Lab, including Dr. Merissa Brousseau, Ryan Klevens, Dr. Andrea Wegrzynowicz, Dr. Vilius Kurauskas, Ashley Hiett, and Levy Treinen for generously sharing their time and expertise discussing membrane protein and multidrug transporter biology. *S. aureus* used as template to obtain the norA gene was a gift from Dr. Helen Blackwell. Some sequencing utilized the University of Wisconsin–Madison Biotechnology Center’s DNA Sequencing Facility (Research Resource Identifier RRID:SCR_017759). RT-qPCR data were obtained at the University of Wisconsin–Madison Biophysics Instrumentation Facility, which was established with support from the University of Wisconsin–Madison and grants BIR-9512577 (NSF) and S10RR013790 (NIH).

## Author contributions

Conceptualization, S.T.M., K.A.H.W., and S.R.; Data curation, S.T.M.; Formal analysis, S.T.M.; Methodology, S.T.M.; Software, S.T.M.; Investigation, S.T.M.; Visualization, S.T.M.; Writing— original draft, S.T.M.; Writing—review and editing, S.T.M., K.A.H.W., and S.R.; Resources, K.A.H.W. and S.R.; Supervision, K.A.H.W. and S.R.; Funding acquisition, K.A.H.W. and S.R.

## Declaration of interests

The authors declare no conflicts of interest.

## Notes

### Competing Interest Statement

The authors have declared no competing interest.

### Summary of Updates

Updated to include author ORCiDs, contributions, and supplemental files

## References

1. Mansoori, B., Mohammadi, A., Davudian, S., Shirjang, S. & Baradaran, B. The Different Mechanisms of Cancer Drug Resistance: A Brief Review. Advanced Pharmaceutical Bulletin 7, 339–348 (2017).

2. Darby, E. M. et al. Molecular mechanisms of antibiotic resistance revisited. Nat Rev Microbiol 21, 280–295 (2023).

3. Neyfakh, A. A. Mystery of multidrug transporters: the answer can be simple. Molecular Microbiology 44, 1123–1130 (2002).

4. Brawley, D. N. et al. Structural basis for inhibition of the drug efflux pump NorA from Staphylococcus aureus. Nat Chem Biol 18, 706–712 (2022).

5. Dou, T., Lian, T., Shu, S., He, Y. & Jiang, J. The substrate and inhibitor binding mechanism of polyspecific transporter OAT1 revealed by high-resolution cryo-EM. Nat Struct Mol Biol 30, 1794–1805 (2023).

6. Minhas, G. S. & Newstead, S. Structural basis for prodrug recognition by the SLC15 family of proton-coupled peptide transporters. Proceedings of the National Academy of Sciences 116, 804–809 (2019).

7. Suo, Y. et al. Molecular basis of polyspecific drug and xenobiotic recognition by OCT1 and OCT2. Nat Struct Mol Biol 30, 1001–1011 (2023).

8. Li, D. C., Nichols, C. G. & Sala-Rabanal, M. Role of a Hydrophobic Pocket in Polyamine Interactions with the Polyspecific Organic Cation Transporter OCT3. J Biol Chem 290, 27633–27643 (2015).

9. Meier, G. et al. Deep mutational scan of a drug efflux pump reveals its structure–function landscape. Nat Chem Biol 19, 440–450 (2023).

10. Wu, H.-H., Symersky, J. & Lu, M. Structure of an engineered multidrug transporter MdfA reveals the molecular basis for substrate recognition. Commun Biol 2, 1–12 (2019).

11. Brill, S., Sade-Falk, O., Elbaz-Alon, Y. & Schuldiner, S. Specificity Determinants in Small Multidrug Transporters. Journal of Molecular Biology 427, 468–477 (2015).

12. Wu, C. et al. Identification of an Alternating-Access Dynamics Mutant of EmrE with Impaired Transport. Journal of Molecular Biology 431, 2777–2789 (2019).

13. Moni, B. M., Quaye, J. A. & Gadda, G. Mutation of a distal gating residue modulates NADH binding in NADH:Quinone oxidoreductase from Pseudomonas aeruginosa PAO1. J Biol Chem 299, 103044 (2023).

14. Klenotic, P. A., Moseng, M. A., Morgan, C. E. & Yu, E. W. Structural and Functional Diversity of Resistance-Nodulation-Cell Division Transporters. Chem Rev 121, 5378–5416 (2021).

15. Khunweeraphong, N., Szöllősi, D., Stockner, T. & Kuchler, K. The ABCG2 multidrug transporter is a pump gated by a valve and an extracellular lid. Nat Commun 10, 5433 (2019).

16. Burata, O. E. et al. Peripheral positions encode transport specificity in the small multidrug resistance exporters. Proc Natl Acad Sci U S A 121, e2403273121.

17. Yee, S. W. et al. The full spectrum of SLC22 OCT1 mutations illuminates the bridge between drug transporter biophysics and pharmacogenomics. Molecular Cell 84, 1932–1947.e10 (2024).

18. Lewinson, O. et al. The Escherichia coli multidrug transporter MdfA catalyzes both electrogenic and electroneutral transport reactions. Proc Natl Acad Sci U S A 100, 1667– 1672 (2003).

19. Tirosh, O. et al. Manipulating the drug/proton antiport stoichiometry of the secondary multidrug transporter MdfA. Proceedings of the National Academy of Sciences 109, 12473– 12478 (2012).

20. Rotem, D. & Schuldiner, S. EmrE, a Multidrug Transporter from Escherichia coli, Transports Monovalent and Divalent Substrates with the Same Stoichiometry*. Journal of Biological Chemistry 279, 48787–48793 (2004).

21. Mazurkiewicz, P., Driessen, A. J. M. & Konings, W. N. Energetics of wild-type and mutant multidrug resistance secondary transporter LmrP of Lactococcus lactis. Biochimica et Biophysica Acta (BBA) – Bioenergetics 1658, 252–261 (2004).

22. Nguitragool, W. & Miller, C. Uncoupling of a CLC Cl−/H+ Exchange Transporter by Polyatomic Anions. Journal of Molecular Biology 362, 682–690 (2006).

23. Grabe, M., Zuckerman, D. M. & Rosenberg, J. M. EmrE reminds us to expect the unexpected in membrane transport. J Gen Physiol 152, e201912467 (2019).

24. Fluman, N., Adler, J., Rotenberg, S. A., Brown, M. H. & Bibi, E. Export of a single drug molecule in two transport cycles by a multidrug efflux pump. Nat Commun 5, 4615 (2014).

25. Wegrzynowicz, A. K. et al. Substrate dependence of transport coupling and phenotype of a small multidrug resistance transporter in Pseudomonas aeruginosa. Journal of Bacteriology 206, e00151–24 (2024).

26. Krupka, R. Channelling free energy into work in biological processes. Experimental Physiology 83, 243–251 (1998).

27. Krupka, R. M. Limits on the Tightness of Coupling in Active Transport. J. Membrane Biol. 167, 35–41 (1999).

28. Krupka, R. M. Uncoupled Active Transport Mechanisms Accounting for Low Selectivity in Multidrug Carriers: P-Glycoprotein and SMR Antiporters. J. Membrane Biol. 172, 129–143 (1999).

29. Jang, S. Multidrug efflux pumps in Staphylococcus aureus and their clinical implications. J Microbiol. 54, 1–8 (2016).

30. Yu, J.-L., Grinius, L. & Hooper, D. C. NorA Functions as a Multidrug Efflux Protein in both Cytoplasmic Membrane Vesicles and Reconstituted Proteoliposomes. J Bacteriol 184, 1370–1377 (2002).

31. Yamada, H. et al. Quinolone susceptibility of norA-disrupted Staphylococcus aureus. Antimicrobial Agents and Chemotherapy 41, 2308–2309 (1997).

32. Papkou, A., Hedge, J., Kapel, N., Young, B. & MacLean, R. C. Efflux pump activity potentiates the evolution of antibiotic resistance across S. aureus isolates. Nat Commun 11, 3970 (2020).

33. Murray, C. J. L. et al. Global burden of bacterial antimicrobial resistance in 2019: a systematic analysis. The Lancet 399, 629–655 (2022).

34. Li, J. et al. Proton-coupled transport mechanism of the efflux pump NorA. Nat Commun 15, 4494 (2024).

35. Xie, P. et al. A fiducial-assisted strategy compatible with resolving small MFS transporter structures in multiple conformations using cryo-EM. Nat Commun 16, 7 (2025).

36. Drew, D., North, R. A., Nagarathinam, K. & Tanabe, M. Structures and General Transport Mechanisms by the Major Facilitator Superfamily (MFS). Chem. Rev. 121, 5289–5335 (2021).

37. Sahin-Tóth, M. & Kaback, H. R. Arg-302 facilitates deprotonation of Glu-325 in the transport mechanism of the lactose permease from Escherichia coli. Proceedings of the National Academy of Sciences 98, 6068–6073 (2001).

38. Starita, L. M. & Fields, S. Deep Mutational Scanning: A Highly Parallel Method to Measure the Effects of Mutation on Protein Function. Cold Spring Harb Protoc 2015, pdb.top077503 (2015).

39. Teelucksingh, T. et al. A genetic platform to investigate the functions of bacterial drug efflux pumps. Nat Chem Biol 18, 1399–1409 (2022).

40. Sulavik, M. C. et al. Antibiotic Susceptibility Profiles of Escherichia coli Strains Lacking Multidrug Efflux Pump Genes. Antimicrob Agents Chemother 45, 1126–1136 (2001).

41. Okusu, H., Ma, D. & Nikaido, H. AcrAB efflux pump plays a major role in the antibiotic resistance phenotype of Escherichia coli multiple-antibiotic-resistance (Mar) mutants. Journal of Bacteriology 178, 306–308 (1996).

42. Nishino, K. & Yamaguchi, A. Analysis of a Complete Library of Putative Drug Transporter Genes in Escherichia coli. J Bacteriol 183, 5803–5812 (2001).

43. Paulsen, I. T. & Skurray, R. A. Topology, structure and evolution of two families of proteins involved in antibiotic and antiseptic resistance in eukaryotes and prokaryotes — an analysis. Gene 124, 1–11 (1993).

44. Griffith, J. K. et al. Membrane transport proteins: implications of sequence comparisons. Current Opinion in Cell Biology 4, 684–695 (1992).

45. Shang, Y. et al. Allele-based analysis revealed the critical functions of region 277–297 in the NorA efflux pump of Staphylococcus aureus. Journal of Antimicrobial Chemotherapy 76, 1420–1427 (2021).

46. Putman, M., van Veen, H. W. & Konings, W. N. Molecular Properties of Bacterial Multidrug Transporters. Microbiol Mol Biol Rev 64, 672–693 (2000).

47. Eilers, M., Patel, A. B., Liu, W. & Smith, S. O. Comparison of Helix Interactions in Membrane and Soluble α-Bundle Proteins. Biophysical Journal 82, 2720–2736 (2002).

48. Henikoff, S. & Henikoff, J. G. Amino acid substitution matrices from protein blocks. Proceedings of the National Academy of Sciences 89, 10915–10919 (1992).

49. Yariv, B. et al. Using evolutionary data to make sense of macromolecules with a “face-lifted” ConSurf. Protein Science 32, e4582 (2023).

50. Tiemann, J. K. S., Zschach, H., Lindorff-Larsen, K. & Stein, A. Interpreting the molecular mechanisms of disease variants in human transmembrane proteins. Biophysical Journal 122, 2176–2191 (2023).

51. Zomot, E. et al. A New Critical Conformational Determinant of Multidrug Efflux by an MFS Transporter. Journal of Molecular Biology 430, 1368–1385 (2018).

52. Cordes, F. S., Bright, J. N. & Sansom, M. S. P. Proline-induced Distortions of Transmembrane Helices. Journal of Molecular Biology 323, 951–960 (2002).

53. Senes, A., Engel, D. E. & DeGrado, W. F. Folding of helical membrane proteins: the role of polar, GxxxG-like and proline motifs. Current Opinion in Structural Biology 14, 465–479 (2004).

54. Yaffe, D., Radestock, S., Shuster, Y., Forrest, L. R. & Schuldiner, S. Identification of molecular hinge points mediating alternating access in the vesicular monoamine transporter VMAT2. Proc Natl Acad Sci U S A 110, E1332–E1341 (2013).

55. Santos, M., Santos, R., Soeiro, P., Silvestre, S. & Ferreira, S. Resveratrol as an Inhibitor of the NorA Efflux Pump and Resistance Modulator in Staphylococcus aureus. Antibiotics (Basel) 12, 1168 (2023).

56. Bhaskar, B. V., Babu, T. M. C., Reddy, N. V. & Rajendra, W. Homology modeling, molecular dynamics, and virtual screening of NorA efflux pump inhibitors of Staphylococcus aureus. Drug Des Devel Ther 10, 3237–3252 (2016).

57. Rodrigues, D. F. et al. Modulation of Drug Resistance by Furanochromones in NorA Overexpressing Staphylococcus Aureus. Evid Based Complement Alternat Med 2022, 9244500 (2022).

58. Palazzotti, D. et al. Fighting Antimicrobial Resistance: Insights on How the Staphylococcus aureus NorA Efflux Pump Recognizes 2-Phenylquinoline Inhibitors by Supervised Molecular Dynamics (SuMD) and Molecular Docking Simulations. J Chem Inf Model 63, 4875–4887 (2023).

59. Morrison, E. A. & Henzler-Wildman, K. A. Transported Substrate Determines Exchange Rate in the Multidrug Resistance Transporter EmrE. J Biol Chem 289, 6825–6836 (2014).

60. Janaszkiewicz, A. et al. Substrate binding and lipid-mediated allostery in the human organic anion transporter 1 at the atomic-scale. Biomedicine & Pharmacotherapy 160, 114342 (2023).

61. Martens, C. et al. Direct protein-lipid interactions shape the conformational landscape of secondary transporters. Nat Commun 9, 4151 (2018).

62. Baker, J. A., Wong, W.-C., Eisenhaber, B., Warwicker, J. & Eisenhaber, F. Charged residues next to transmembrane regions revisited: “Positive-inside rule” is complemented by the “negative inside depletion/outside enrichment rule”. BMC Biology 15, 66 (2017).

63. Zhang, X. C., Zhao, Y., Heng, J. & Jiang, D. Energy coupling mechanisms of MFS transporters. Protein Sci 24, 1560–1579 (2015).

64. Klyachko, K. A., Schuldiner, S. & Neyfakh, A. A. Mutations affecting substrate specificity of the Bacillus subtilis multidrug transporter Bmr. Journal of Bacteriology 179, 2189–2193 (1997).

65. Smirnova, I., Kasho, V., Sugihara, J., Choe, J.-Y. & Kaback, H. R. Residues in the H+ Translocation Site Define the pKa for Sugar Binding to LacY. Biochemistry 48, 8852–8860 (2009).

66. Schuldiner, S. Competition as a Way of Life for H+-Coupled Antiporters. Journal of Molecular Biology 426, 2539–2546 (2014).

67. Zwama, M. & Yamaguchi, A. Molecular mechanisms of AcrB-mediated multidrug export. Research in Microbiology 169, 372–383 (2018).

68. Bodosa, J. & Klauda, J. B. Metadynamics Study of Lipid-Mediated Antibacterial Toxin Binding to the EmrE Multiefflux Protein. J. Phys. Chem. B 128, 8712–8723 (2024).

69. Adler, J., Lewinson, O. & Bibi, E. Role of a conserved membrane-embedded acidic residue in the multidrug transporter MdfA. Biochemistry 43, 518–525 (2004).

70. Golin, J. et al. Studies with Novel Pdr5p Substrates Demonstrate a Strong Size Dependence for Xenobiotic Efflux*. Journal of Biological Chemistry 278, 5963–5969 (2003).

71. Varela, M. F., Sansom, C. E. & Griffith, J. K. Mutational analysis and molecular modelling of an amino acid sequence motif conserved in antiporters but not symporters in a transporter superfamily. Molecular Membrane Biology 12, 313–319 (1995).

72. Zhang, Y. et al. Understanding Cytochrome P450 Enzyme Substrate Inhibition and Prospects for Elimination Strategies. ChemBioChem 25, e202400297 (2024).

73. Stein, K. C. & Frydman, J. The stop-and-go traffic regulating protein biogenesis: How translation kinetics controls proteostasis. Journal of Biological Chemistry 294, 2076–2084 (2019).

74. Ng, E. Y., Trucksis, M. & Hooper, D. C. Quinolone resistance mediated by norA: physiologic characterization and relationship to flqB, a quinolone resistance locus on the Staphylococcus aureus chromosome. Antimicrob Agents Chemother 38, 1345–1355 (1994).

75. Slonczewski, J. L., Fujisawa, M., Dopson, M. & Krulwich, T. A. Cytoplasmic pH Measurement and Homeostasis in Bacteria and Archaea. in Advances in Microbial Physiology (ed. Poole, R. K.) vol. 55 1–317 (Academic Press, 2009).

76. Booth, I. R. Regulation of cytoplasmic pH in bacteria. Microbiological Reviews 49, 359–378 (1985).

77. Padan, E., Bibi, E., Ito, M. & Krulwich, T. A. Alkaline pH homeostasis in bacteria: New insights. Biochimica et Biophysica Acta (BBA) – Biomembranes 1717, 67–88 (2005).

78. Padan, E., Zilberstein, D. & Schuldiner, S. pH homesstasis in bacteria. Biochimica et Biophysica Acta (BBA) – Reviews on Biomembranes 650, 151–166 (1981).

79. Slonczewski, J. L., Rosen, B. P., Alger, J. R. & Macnab, R. M. pH homeostasis in Escherichia coli: measurement by 31P nuclear magnetic resonance of methylphosphonate and phosphate. Proc Natl Acad Sci U S A 78, 6271–6275 (1981).

80. Jiang, D. et al. Structure of the YajR transporter suggests a transport mechanism based on the conserved motif A. Proc. Natl. Acad. Sci. U.S.A. 110, 14664–14669 (2013).

81. Brousseau, M., Teng, D., Thomas, N. E., Voth, G. A. & Henzler-Wildman, K. A. The C-terminus of the multi-drug efflux pump EmrE prevents proton leak by gating transport. Preprint at 10.7554/eLife.105525.1 (2025).

82. Miller, S. T., MacDonald, C. B. & Raman, S. Understanding, inhibiting, and engineering membrane transporters with high-throughput mutational screens. Cell Chemical Biology 32, (2025).

83. Doshi, R., Nguyen, T. & Chang, G. Transporter-mediated biofuel secretion. Proceedings of the National Academy of Sciences 110, 7642–7647 (2013).

84. Kell, D. B., Swainston, N., Pir, P. & Oliver, S. G. Membrane transporter engineering in industrial biotechnology and whole cell biocatalysis. Trends Biotechnol 33, 237–246 (2015).

85. Piotrowski, J. et al. Death by a thousand cuts: the challenges and diverse landscape of lignocellulosic hydrolysate inhibitors. Frontiers in Microbiology 5, (2014).

86. Zhang, L. et al. Bacterial Efflux Pump Inhibitors Reduce Antibiotic Resistance. Pharmaceutics 16, 170 (2024).

87. Wegrzynowicz, A., Spreacker, P., Demas, S., Powers, E. & Henzler-Wildman, K. Inducing susceptibility with a Small Multidrug Resistance transporter from P. aeruginosa. Physiology 38, 5787819 (2023).

88. Spreacker, P. J. et al. Activating alternative transport modes in a multidrug resistance efflux pump to confer chemical susceptibility. Nat Commun 13, 7655 (2022).

89. Tetko, I. V. et al. Virtual computational chemistry laboratory--design and description. J Comput Aided Mol Des 19, 453–463 (2005).

90. Knox, C. et al. DrugBank 6.0: the DrugBank Knowledgebase for 2024. Nucleic Acids Research 52, D1265–D1275 (2024).

91. Kim, S. et al. PubChem 2025 update. Nucleic Acids Research 53, D1516–D1525 (2025).

92. Goodman, D. B., Church, G. M. & Kosuri, S. Causes and effects of N-terminal codon bias in bacterial genes. Science 342, 475–479 (2013).

93. Datsenko, K. A. & Wanner, B. L. One-step inactivation of chromosomal genes in Escherichia coli K-12 using PCR products. Proceedings of the National Academy of Sciences 97, 6640– 6645 (2000).

94. Baba, T. et al. Construction of Escherichia coli K-12 in-frame, single-gene knockout mutants: the Keio collection. Mol Syst Biol 2, 2006.0008 (2006).

95. Scholz, S. A. et al. High-Resolution Mapping of the Escherichia coli Chromosome Reveals Positions of High and Low Transcription. Cell Syst 8, 212–225.e9 (2019).

96. Wiegand, I., Hilpert, K. & Hancock, R. E. W. Agar and broth dilution methods to determine the minimal inhibitory concentration (MIC) of antimicrobial substances. Nat Protoc 3, 163– 175 (2008).

97. Zhang, J., Kobert, K., Flouri, T. & Stamatakis, A. PEAR: a fast and accurate Illumina Paired-End reAd mergeR. Bioinformatics 30, 614–620 (2014).

98. Rubin, A. F. et al. A statistical framework for analyzing deep mutational scanning data. Genome Biology 18, 150 (2017).

99. Zhou, K. et al. Novel reference genes for quantifying transcriptional responses of Escherichia coli to protein overexpression by quantitative PCR. BMC Mol Biol 12, 18 (2011).

100. Lomize, M. A., Pogozheva, I. D., Joo, H., Mosberg, H. I. & Lomize, A. L. OPM database and PPM web server: resources for positioning of proteins in membranes. Nucleic Acids Res 40, D370–376 (2012).

101. Koehler Leman, J., Mueller, B. K. & Gray, J. J. Expanding the toolkit for membrane protein modeling in Rosetta. Bioinformatics 33, 754–756 (2017).

102. Alford, R. F. et al. An Integrated Framework Advancing Membrane Protein Modeling and Design. PLoS Comput Biol 11, e1004398 (2015).

103. Fleishman, S. J. et al. RosettaScripts: A Scripting Language Interface to the Rosetta Macromolecular Modeling Suite. PLOS ONE 6, e20161 (2011).

104. Park, H. et al. Simultaneous Optimization of Biomolecular Energy Functions on Features from Small Molecules and Macromolecules. J. Chem. Theory Comput. 12, 6201–6212 (2016).

